# Chronic Inorganic Fertilization Shifts Evolutionary Trajectories and Induces a Regulatory Shield in Maize Rhizosphere Bacteria

**DOI:** 10.64898/2026.06.04.729967

**Authors:** Bharat Mishra, Anuradha Goswami

## Abstract

Long-term agricultural fertilization is a major driver of structural shifts in the soil microbiome, yet traditional metagenomic analyses often fail to capture the population-level evolutionary dynamics and heritable genetic adaptations that dictate species survival in the rhizosphere. Here, we built and utilize a high-resolution, gene-centric population-genomic framework to resolve the microdiversity and selective landscapes shaping maize rhizosphere bacteria across a chronic management gradient including Inorganic NPK fertilization, Organic compost amendment, and an Unfertilized Control. We observed a distinct decoupling between total mutational volume and adaptive fixation rates. Inorganic fertilization acts as a massive genotoxic hotspot generating 5,544 variants but maintaining a baseline intra-population variants consensus rate of 37.3% due to intense purifying selection. Whereas organic management restricts global mutational volume but drives relatively high rate of 49.6%, signaling a highly coordinated, directional selection where transposable elements function as bridges for structural adaptations. Crucially, we uncover that regulatory shield mechanism that protects the core genome from deleterious mutations while core metabolic and structural genes exhibit absolute sequence stasis. Additionally, we report that mutations were precisely accumulated within the immediate 0–200 bp promoter region of key stress-defense and metabolic genes. This targeted regulatory remodeling is chemically fingerprinted by a depressed transition-to-transversion ratio and structurally supported by Mobile Fortress model that significantly expands the repository of Highly Conserved Mobile Loci sequestered on cryptic circular plasmids. Together, these evolutionary trajectories indicate that long-term inorganic fertilization forces rhizosphere bacteria to prioritize cellular repair and resource hoarding over host interactions. By demonstrating how high-input management shifts selection from cooperative pathways toward internal defense, these findings provide a population-genomic explanation for the suppression of biological nutrient-use efficiency observed in intensive agroecosystems.

## Introduction

Fertilization is the dominant management practice in crop agriculture and, incidentally, one of the largest perturbations humans impose on the soil microbiome. Inorganic, organic, and integrated fertilization regimes shape soil microbial communities through divergent mechanisms with measurable functional consequences (1–3). Long-term inorganic nitrogen application shifts soil pH, microbial community composition, biomass carbon, and enzymatic activities involved in nitrogen and phosphorus cycling (4, 5). Whereas organic fertilizers including manure, compost, cover-crop residues produce the inverse signature by increases microbial biomass, diversify functional gene repertoires across nitrogen-cycling, carbon-decomposition guilds, and stimulate fungal-to-bacterial biomass ratios (5, 6). Furthermore, the soil resistome shape the mobilization of accessory and resistance genes through horizontal gene transfer (7). Traditional investigations of soil microbial communities have primarily focused on compositional variance and gene abundances (8), which often overlooks the exploration of heritable traits that determine membership within individual microbial species. Connecting the observed community back to genetic traits will be significant next step in population-genomic of genome-resolved metagenomics.

Long-term inorganic fertilization and modern maize breeding under high-nitrogen conditions are profoundly restructuring the rhizosphere microbiome, shifting it away from ancestral microbial dependencies (9, 10). Continual reliance on synthetic fertilizers has successfully maximized global grain yields, but it has simultaneously suppressed the plant’s natural recruitment of beneficial diazotrophs, the specific bacteria responsible for biological nitrogen fixation (11). Because modern germplasm has been selected and developed in nutrient-saturated environments, these domesticated maize varieties no longer invest energy into attracting nitrogen-fixing partners. Instead, their root zones assemble specialized microbial communities characterized by a larger biosynthetic gene cluster. This evolutionary trade-off effectively replaces a self-sustaining, microbially driven nutrient cycle with an artificial system heavily dependent on continuous chemical inputs and industrial soil management (12, 13). While this fertilizer influenced rhizosphere taxonomic changes are now studied at metagenomic level, the fundamental genomic feature changes underlying the microbial adaptation to chemical stressors remain largely unexplored.

The genomic basis of microbial adaptation to fertilization in soil has been limited by the relevant and measurable signal, mutagenic potential of nitrogen metabolism, and integrated host regulatory response. With the recent democratization of high throughput shotgun metagenomes and availability of high quality metagenomic assemblies and analysis tools can reveal the genomic features including within-population nucleotide diversity, selection pressure, fixation indices, and recombination(14, 15). Recently, these approaches have been used to quantify gene-specific selection in grassland soil (16) and in soil-dwelling pangenomes (17), but not across a fertilization gradient. Tools and approaches to identify the chromosomal and non-chromosomal mutations including plasmids and mobile genetic elements in fertilized soil metagenome are very limited. To address this technical and knowledge gap, the present study quantifies evolutionary velocity within maize rhizosphere metagenomes, offering insight into the processes by which microbial populations adapt to fertilization regimes. We hypothesize that chronic inorganic/synthetic chemical stress restricts mutation of certain essential genes in the microbial community which are conserved and crucial for survival of the community. To test this hypothesis, we built a reproducible framework *MAGmutome* and conducted a high-resolution discovery-based metagenomic survey to characterize the landscape of genetic variation across three distinct rhizosphere populations. Additionally, our genomics-integrated evolutionary study provides a high-resolution perspective on multi-factorial inorganic fertilizer management (MFIFM) with long-term extensive and intensive chemically stressed environment during with cultivated and ploughing farming practice in maize root and bulk soil microbes. Further, the findings of this study demonstrate that microbial survival under chemical stress is facilitated by an evolutionary stratification within the metagenome. Moreover, the fertilizer impact on the root transcriptomics also demonstrate that stress response and nutrient transport pathways are more impacted in maize. Overall, the high-resolution genomic analysis of adaptive responses under different agricultural practices highlights the mechanisms through which microbial populations function under stress, contributing to a deeper understanding of microbial resilience in agricultural soils.

## Results and Discussion

### In situ selection coefficients and adaptive flux reveal microbial survival strategies under inorganic chemical stress

The long-term (chronic) fertilizer application is known for disturbances in the soil microbial community and tilts community composition toward fast-growing copiotrophs at the expense of slow-growing oligotrophs (18). Long-term application of both organic compost and inorganic chemical fertilizers drastically alters the soil microbiome by reducing overall evenness. High nutrient inputs favor fast-growing copiotroph genera (*Bacillus, Pseudomonas, Burkholderia*), allowing them to outcompete and displace the diverse oligotrophic baseline community (*Streptomyces, Mycobacterium, Methylobacterium, Bradyrhizobium*) found in the control soil (Fig. 1a). This effect is most pronounced in the inorganic fertilizer treatment. Additional analysis of the multi-factorial inorganic fertilizer management (MFIFM) regimes preserves a highly diverse and structurally stable microbial matrix (Fig. S1a). While tillage and nitrogen intensity introduce minor profile changes, the Rhizosphere effect remains the primary driver of community variation. It selectively enriches fast responding copiotroph genera (specifically *Pseudomonas*) due to the abundance of root exudates, while the broader, highly structured oligotrophic baseline (led by *Streptomyces*, *Mycobacterium*) provides the background functional buffering capacity of the soil ecosystem.

**Figure 1.**
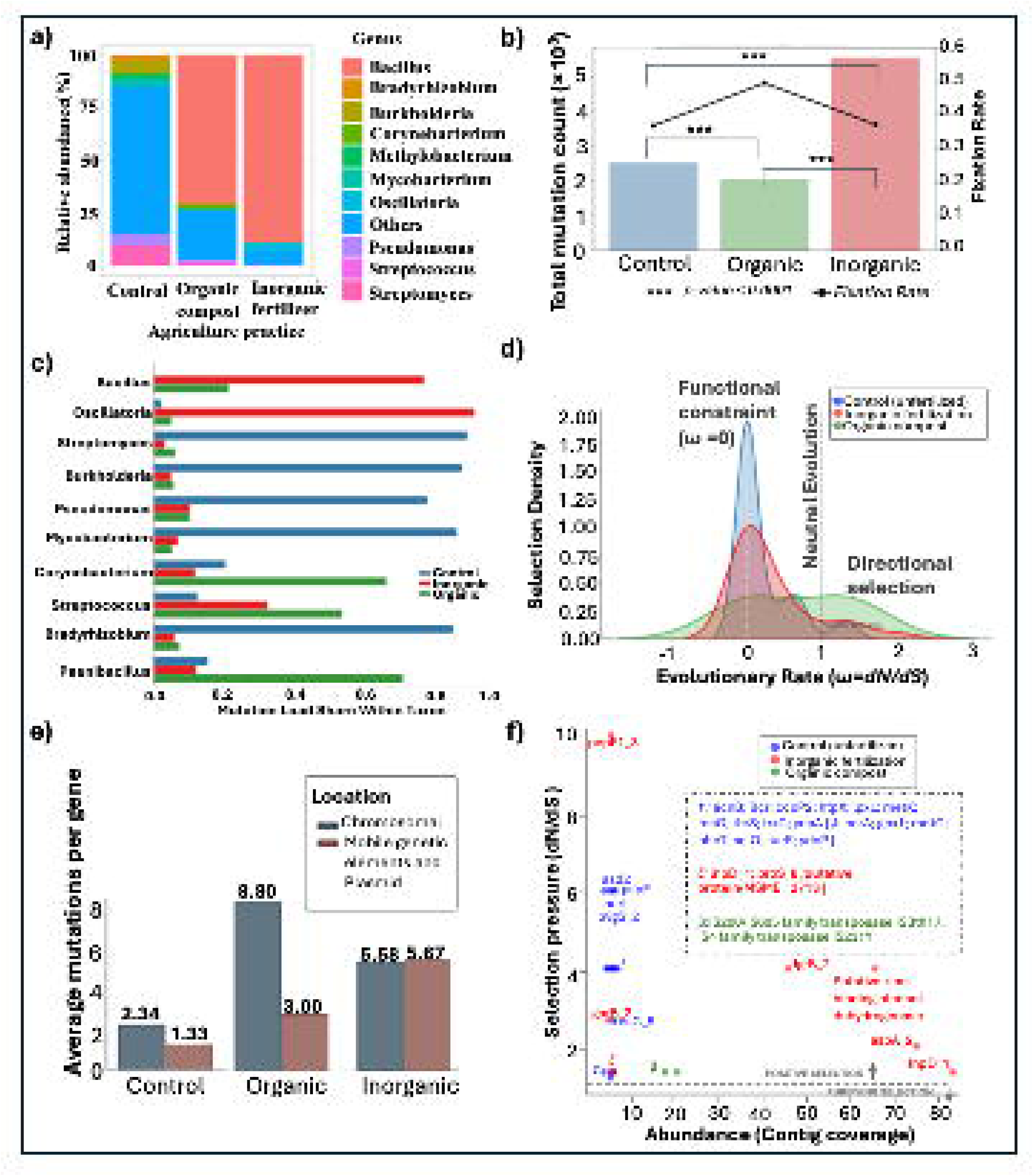
Microbial selection and adaptive survival strategies under fertilizer regimen. **(a).** The microbial genus taxonomic relative abundance plot in different agricultural practices. Only top 10 taxa are displayed. **(b)**. The dual-axis plot displays absolute mutational burden (bars) against the population fixation rate (AF = 1.0; black line). Inorganic fertilization (red) demonstrates high absolute mutation counts but baseline fixation whereas Organic management (green) demonstrates a background mutation but significantly higher fixation rates (∼50%, *p-value <0.0001*). **(c)**. The mutation load share within microbial taxon at genus level with each fertilizer treatment strategy (Blue: control, green: organic: red: inorganic). **(d)**. The selection density represents the probability distribution of evolutionary rates measured as in-situ selection coefficient (dN/dS) across a genomic dataset. **(e)**. The average number of mutations in per chromosomal and non-chromosomal genes in three fertilizer treatments. **(f)**. Selection pressure of genes with either purifying or positive selection and the contig coverage in fertilizer treatments.

To investigate how long-term agricultural practices shape the genomic evolution of soil microbial populations, we quantified genetic heterogeneity and selective pressure across three regimes including Inorganic fertilization, Organic compost, and a non-fertilized Control. The Single Nucleotide Polymorphisms (SNPs) analysis identified a total of 7,582 high-confidence variant across the three populations (Fig. 1b, Fig S2a). The inorganic metagenome exhibited a significantly higher density of genetic flux (n = 5,544 variants) compared to the organic population (n = 2,038; Mann-Whitney U, p < 0.001). However, despite this mutation volume, the proportion of population-consensus variants or Fixation rate (defined as alleles reaching an Allele Frequency of 1.0 within the sampled library) was 37.3%, which was equivalent to the baseline control (36.9%). Conversely, the organic management regime demonstrated a distinct genomic trajectory despite a more moderate mutation load (2,038 SNPs), it achieved the highest fixation state at 49.6% (Fig. 1b). Elevated mutation counts and fixation rates were also distinctly noted within the intensive and cultivator samples of the MFIFM cohort. (Fig. S1b).

To identify the impact of fertilizer regimen mutation across genus, we explored the mutation load within different taxon. The distribution of the mutation load share within the top 10 genus microbial community exhibited a distinct, taxon-specific dependency on fertilizer treatment conditions (Fig. 1c, Fig S2b). Most of the genera specifically *Oscillatoria* and *Bacillus* mutation load share is driven by the Inorganic treatment, reaching approximately 0.90 and 0.75, respectively. Other genera including *Streptomyces*, *Burkholderia*, *Pseudomonas*, *Mycobacterium*, and *Bradyrhizobium* mutation load share is heavily localized within Control condition, with each exceeding a relative proportion of *0.80*, while remaining minimal (*< 0.10*) under both fertilizer treatments. Interestingly, we identified a distinct response in *Corynebacterium*, *Streptococcus*, and *Paenibacillus* with elevated susceptibility or evolutionary acceleration primarily under the Organic fertilizer treatment, yielding localized mutation shares ranging between 0.50 and 0.70. Finally, the MFIFM cohort demonstrated a highly uniform macro-pattern across the community, where intensive environments systematically acted as the primary evolutionary drivers, maximizing relative mutation shares between 0.15 and 0.19 across nearly all surveyed genera (Fig. S2c-d). Collectively, these data indicate a robust interaction effect between bacterial phylogeny and environmental selective pressures, demonstrating that xenobiotic or nutritional stressors do not uniformly elevate mutation rates across a community, but instead induce highly localized genomic impacts restricted to specific target taxa (19, 20).

To explore the mutational burden on microbial genes and selective pressure, first we calculated the total number of genes affected by mutations in different fertilizer regimen (Table 1). The control sample exhibited the highest overall genetic diversity, with 872 mutations distributed across 375 unique genes. Conversely, both organic (50 mutations) and inorganic (335 mutations) fertilization regimes significantly restricted the number of genes affected narrowing to 7 and 60 genes, respectively. Despite this narrowing, the intensity of selection per gene was notably higher under fertilization. The organic sample exhibited an average of 7.14 mutations per gene, nearly triple the rate observed in the control group (2.33 mutations/gene) while inorganic fertilization exhibits an average of 5.58 mutations per gene. Additionally, analysis of mutational velocity (Vpkb) confirmed a marginal trend toward global variation (Fig. S3a; Kruskal-Wallis H = 5.36, p = 0.068). Pairwise investigations revealed that organic amendment significantly shifted the mutational landscape compared to the control (Fig. S3a; Mann-Whitney U, p = 0.0273), supported by a large effect size (Cliff’s Delta d = −0.487). This suggests that organic stress creates a distinct selective signal rather than a stochastic occurrence. In contrast, inorganic fertilization did not significantly alter overall mutation density relative to the control (p = 0.3706, d = 0.072), indicating that NPK stress, while altering taxonomic abundance does not induce the same density of mutational hotspots as organic practices (10). <u>The MFIFM cohort treatments shared similar spread among</u> <u>different inorganic fertilization and agricultural practices (Fig. S3b).</u>

**Table 1.** A summary statistics of mutation volume per gene impacted by long-term agricultural practices including different fertilization regimes.

| *Condition* | *Total Mutations* | *Genes affected (n)* | *Median variants per kilobase (Vpkb)* | *Average mutation per gene* | *Mann-Whitney U test Significance (vs Control)* |
| --- | --- | --- | --- | --- | --- |
| **Control** | 872 | 375 | 0.293 | 2.33 | *Reference* |
| **Inorganic** | 335 | 60 | 0.288 | 5.58 | P = 0.37; d = 0.07 (*Negligible*) |
| **Organic** | 50 | 7 | 0.087 | 7.14 | P = 0.03; d = -0.49 (*Large*) |

Next, we examined the selection density and identified a wide, low-density distribution centered between 0.75 and 1.0 across all treatments including control, organic, and inorganic (Fig. 1d). Control and inorganic soil treatments show a sharp rise in density at *dN/dS* = 0, suggesting that synonymous mutations accumulate while harmful non-synonymous mutations are eliminated by natural selection. By contrast, when *dN/dS* > 1, organic treatment seems to promote directional selection in certain chromosomal genes. Interestingly, the MFIFM cohort follow similar pattern of selection density across all treatments suggesting that most of harmful non-synonymous mutations are eliminated by natural selection (Fig. S4). Additionally, we explored the mutation rate in different genomic regions and report that chromosomal genes have very high mutations per gene in organic (8.80) compared to inorganic (5.58) and control (2.34) (Fig. 1e). This high intensity was primarily driven by a chromosomal hypothetical protein (38 mutations) and specific metabolic/regulatory genes including carD (transcriptional regulator), queG (epoxyqueuosine reductase), and glgB (glycogen branching enzyme) (Fig. S5). Whereas extra chromosomal genes including mobile genetic elements (MGE) and plasmids mutation rate is higher in the inorganic treated root rhizosphere.

In inorganic conditions, the mutational load was almost equal between chromosomal (5.58) and mobile genetic elements/plasmid (5.67) (Fig. 1e). This might indicate a broad-spectrum stress response where the bacteria evolve their core genomes and their mobile at the same rate. Control conditions depict a stable population undergoing neutral genetic drift than targeted environmental selection.Finally, to distinguish the role of neutral genetic drift and functional adaptation, we correlated the in-situ selection coefficient (*dN/dS*) of high-selection genes with their respective contig abundances (Fig. 1f). We observed distinct selective sweeps under inorganic fertilization, most notably in the pepF1_3 gene (*dN/dS* = 10.0), suggesting a rapid functional shift in response to chemical stress (Fig. 1f). Conversely, the organic compost regime was characterized by positive selection on mobile genetic elements (transposases), indicating that adaptation to organic amendments may be mediated by genomic reorganization and horizontal gene transfer.Overall, these findings demonstrate that while inorganic stress drives specific microbial community changes, mutation count, and functional sweeps, the organic practices promote a broader, potentially more flexible, evolutionary strategy (21).

### Inorganic fertilization practices alter the DNA lesion profile mutational signatures

To explore the mutational spectrum of fertilization regimes to identify the potential drivers of genomic instability we studied the mutational nucleotide landscape of the soil microbiome. We observed a profound shift in the transition-to-transversion ratio compared to the stable population in control (Fig. 2a). Specifically, a highly significant enrichment in transition mutations was identified, with a directional bias toward C→T (p = 6.94 ×10^-10^) and G→A (p = 1.07 ×10^-5^) (Supplementary File). These shifts represent the biochemical hallmarks of oxidative deamination, where inorganic nitrogen salts likely accelerate the conversion of cytosine to thymine. The most extreme divergence was observed in the T→C transition rate (p = 4.76 ×10^-17^), suggesting a systemic failure in DNA mismatch repair or a specific polymerase bias induced by the inorganic environment (22). Furthermore, the Inorganic treatment exhibited a significant spike in transversions, specifically A→T and T→A (p < 10^-10^). Unlike transitions, these transversions are indicative of more severe genotoxic stress, likely mediated by Reactive Oxygen Species (ROS) or ionic imbalances. Overall, the surge in both transitions (∼3,500) and transversions (∼2,100) in microbial population experiencing inorganic fertilization regime show genomic distress conditions (Fig. S6). The MFIFM cohort analysis also revealed similar mutational signatures with higher enrichment in transition mutations (A → G, C → T, G → A, and T → C) in all regimens, whereas the transversions mutations (A→ T and T→A) are more enriched in the bulk soil with cultivator and Intensive fertilization mimicking very high the Inorganic treatment (Fig. S7a-b).

**Figure 2.**
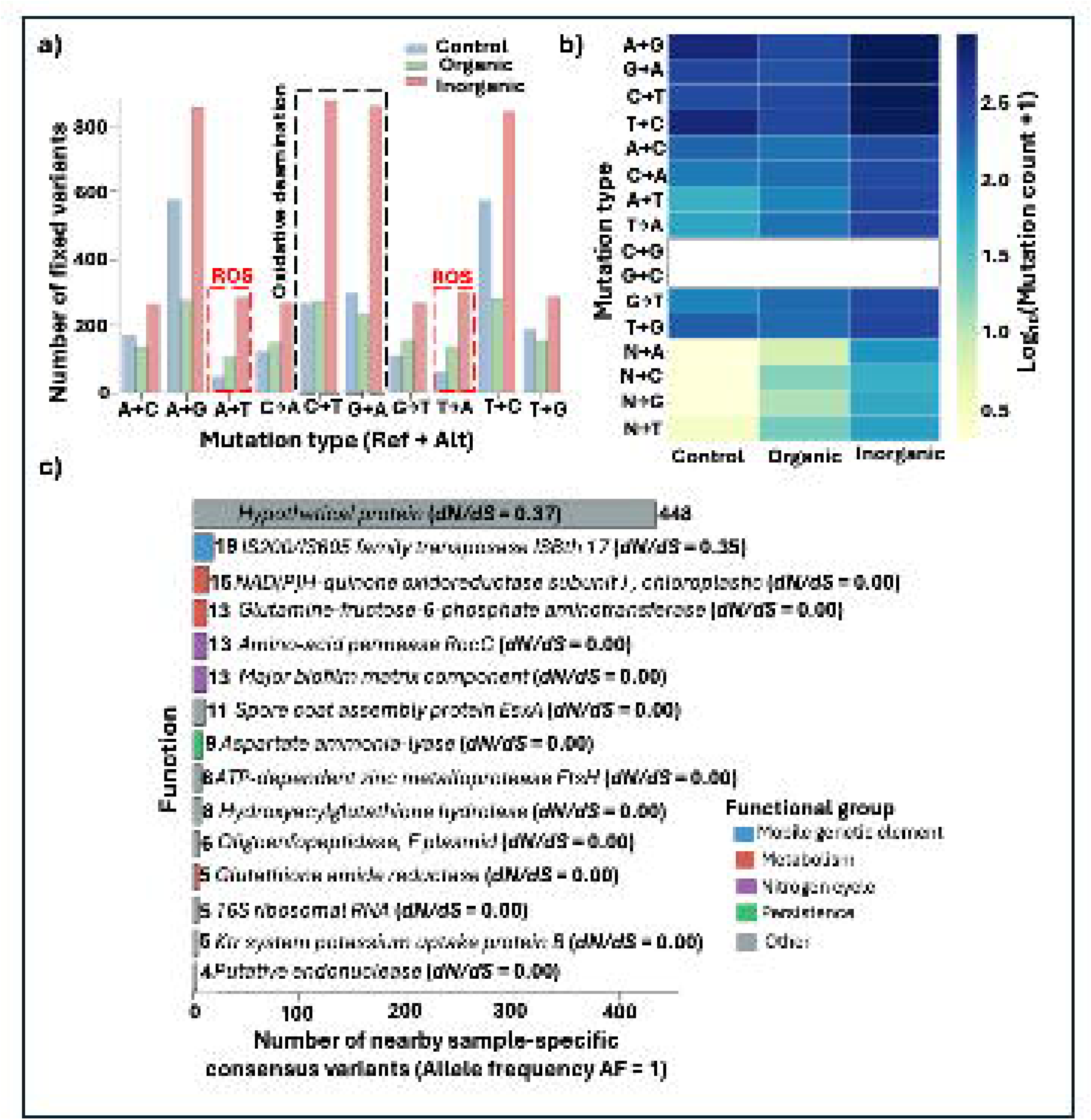
Comparative Mutational Signatures Across Soil Management Regimes. **(a).** Distribution of primary transition and transversion events. Inorganic fertilization induced a systemic mutational surge, characterized by a significantly higher frequency of C →T transitions (p = 6.94 ×10^-10^) and A →T transversions (p = 1.42 ×10^-11^). Notably, the Organic treatment displays a genomic buffering effect, maintaining mutational profiles comparable to the unamended Control. The absence of G ⟷C transversions across all samples underscores the inherent thermodynamic stability of triple-hydrogen-bonded pairs despite environmental stress. **(b).** Mutational intensity heatmap of single-base substitutions (log-transformed counts) across Control, Organic, and Inorganic agricultural management regimes. **(c)**. Functional enrichment of mutated regulatory shield mechanism genes accumulating high frequencies of fixed intergenic mutations within the promoters. Shock hits color-coded by functional category: Nitrogen Metabolism (Green), Genetic Mobility (Blue), and Stress/Secretion (Purple).

The comparative analysis between mutational load and selection intensity revealed a fundamental evolutionary paradox (23). While the inorganic treatment functioned as a mutational hotspot, the inorganic chemical stress did not translate into functional diversification. Instead, core metabolic genes including accC (biotin carboxylase), ade_1, and inorganic triphosphatase—maintained a mutation velocity (*dN/dS*) of 0.0 (Supplementary file with dN/dS data). It suggests that under high inorganic stress, non-synonymous mutations in essential genes leading to genomic stability undergoes purifying selection was further supported by molecular evidence. The mutational profile in inorganic fertilization (Table 2), was dominated by mutation type caused by polymerase errors or oxidative lesions (C → T and A → T) while no mutation observed at G ⟷C transversions (Fig. 2b) revealing that the purifying selection or high-fidelity repair systems were successfully filtering out mutations at these high-stability sites to prevent deleterious changes to core protein functions.

**Table 2.** Comparison of Mutational Profiles Across All Treatments.

| Substitution | Control (Ctrl) | Organic | Inorganic | Organic Fold Change (vs Ctrl) | Inorganic Fold Change (vs Ctrl) | Maximum significance (p) |
| --- | --- | --- | --- | --- | --- | --- |
| $C \rightarrow T$ | 268 | 274 | 879 | 1.02x | 3.28x | $6.94 \times 10^{-10}$ |
| $G \rightarrow A$ | 298 | 234 | 864 | 0.78x | 2.90x | $1.07 \times 10^{-5}$ |
| $A \rightarrow T$ | 48 | 106 | 287 | 2.21x | 5.98x | $1.42 \times 10^{-11}$ |
| $T \rightarrow A$ | 60 | 138 | 304 | 2.30x | 5.06x | $7.61 \times 10^{-10}$ |
| $T \rightarrow C$ | 579 | 283 | 847 | 0.49x | 1.46x | $4.76 \times 10^{-17}$ |
| $A \rightarrow G$ | 581 | 278 | 860 | 0.48x | 1.48x | $2.62 \times 10^{-16}$ |
| $N \rightarrow \text{Any}$ | 5 | 55 | 254 | 11.0x | 50.8x | $1.00 \times 10^{-9}$ |

Furthermore, the comparative analysis of spatial mutational pressure highlighted a distinct disparity in regulatory and coding evolution across treatments. The inorganic fertilization induces a severe genomic shock in rhizosphere microbial communities, driving a distinct regulatory-focused evolutionary strategy that is entirely mitigated by organic soil management (Fig. 2c). Under inorganic nitrogen stress, microbial populations employ a regulatory shield mechanism, accumulating high frequencies of fixed intergenic mutations within the promoters of core metabolic (NAD(P)H-quinone oxidoreductase (15 hits) and Glutamine-fructose-6-phosphate aminotransferase (13 hits)), stress-response, redox, and biofilm-mediating genes (such as *ftsH* proteases and glutathione reductases) while maintaining a dN/dS ratio of 0.0 (Fig. 2c). This indicates intense purifying selection to modulate gene expression without altering critical protein structures. Conversely, the organic treatment exhibits a total absence of these regulatory hotspots, shifting mutational pressure toward background or coding-sequence adaptations (Fig. S8a). This evolutionary divergence is most pronounced in mobile genetic elements like the *IS200/IS605* family transposases (19 hits) because inorganic stress forces strict regulatory control over transposition frequency to maintain genomic stability (Fig. 2c), whereas organic soil management alleviates this pressure, triggering positive selection within the coding sequences (dN/dS = 1.33) to utilize these elements as adaptive bridges for functional diversification (Table S3). The MFIFM cohort analysis also revealed similar regulatory hotspots across treatments (Fig. S8b). These results posit a fundamental shift in strategy, while inorganic fertilization forces the bacteria to strictly regulate the jumping frequency of mobile genetic elements to maintain stability, organic soil allows these elements to act as an adaptive bridge, where the proteins themselves actively evolve to become more efficient tools for functional diversification

### Evolutionary dynamics of inorganic fertilization select for genomic compartmentalization

To investigate the evolution of promoters governing metabolism and other regulation we explored the genomic shock describing stress induced mutagenesis across fertilizer regimen. We observed a total of 5,544 fixed mutations (sample-specific consensus variants (or AF=1.0 variants)) in the inorganic treatment, of which only 347 (6.2%) were intragenic, while a vast majority (5,197 fixed variants) were localized within intergenic regions (Fig. 3a). Proximity analysis revealed that these intergenic mutations were not uniformly distributed. Instead, they exhibited a bifurcated spatial architecture. i) *Orphan Variants (n = 4,835)* are most of the population consensus (fixed variants) occurred in a non-clustered, stochastic distribution across deep intergenic regions. These orphan variants represent the baseline of neutral genetic drift, reflecting the background evolutionary noise of the soil community. The 12 identified functional islands (red dots) exhibit a mutational density in several higher order of magnitude than the genomic mean (Fig. 3a). The next spatial architecture is ii) *Precision Regulatory Hits (n = 705)* are mutations exhibited extreme spatial precision, clustering within the immediate 0 – 200 bp promoter regions of essential metabolic and plasticity genes (Fig. 3b). This concentration of variants within transcriptional initiation sites indicates that the selective pressure of inorganic fertilization primarily focusses on regulation of metabolic and plasticity-related genes rather than altering their protein structures (24).

**Figure 3.**
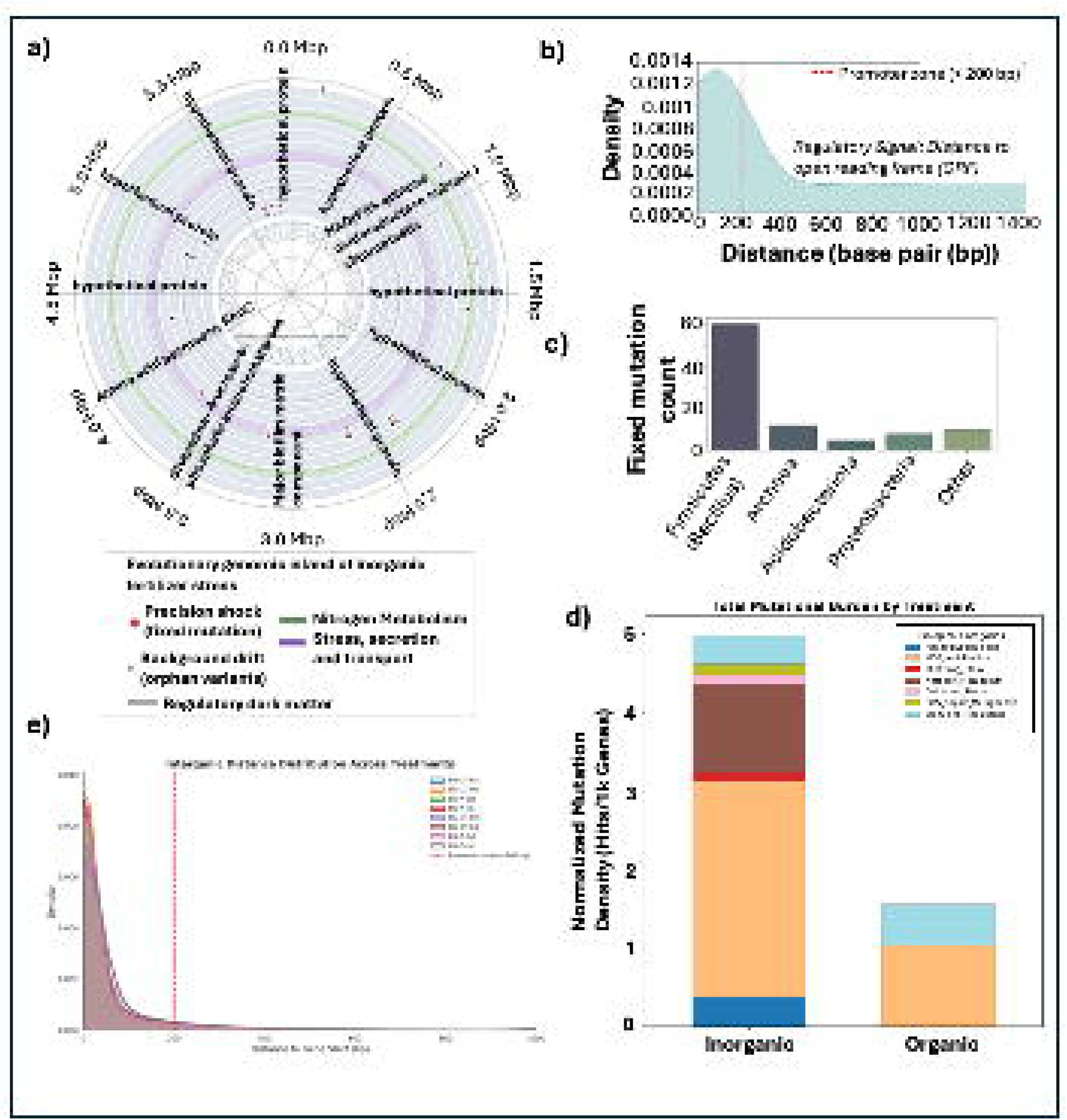
Spatial localization of fixed mutations relative to functional hotspots and taxonomic origin. **(a).** Polar Distribution of Mutational Pressure Circular representation of fixed mutations (AF = 1.0) across the dominant metagenomic scaffolds of the inorganic nitrogen treatment. Precision Shock Tracks (Outer Rings) Individual tracks represent key functional islands showing significant mutational enrichment. Red markers indicate the exact spatial coordinates of fixed mutations within regulatory (intergenic) hotspots. Genomic Drift / Orphan Variants (Inner Cloud): The grey stippled region represents the distribution of *n=4,835* orphan variants. **(b).** Distributions of mutation density (Hits/kb) across genomic compartments under inorganic NPK stress high density within the immediate 0 – 200 bp promoter regions. **(c)**. Taxonomic anchoring and functional mapping of genomic shock in inorganic fertilization. **(d).** The normalized mutation density enrichment in in different biological categories in inorganic and organic fertilizer treatment. (e). The Distributions of mutation density (Hits/kb) across genomic compartments in different treatment groups of MFIFM cohort high density within the immediate 0 – 200 bp promoter regions.

Next, we mapped the taxonomic anchoring and functional mapping of genomic shock in inorganic fertilization. The alignment of our genomic shock regions with functional BLAST data confirms that inorganic fertilization triggers a community-wide selective sweep. While 4,835 orphan variants represent the background genomic drift of this high-velocity environment, the 705 precision hits represent the functional core of survival. This core is dominated by the optimization of nitrogen processing, the activation of mobile genetic elements, and the reinforcement of cellular defense mechanisms across both Bacterial and Archaeal domains (Fig. 3c, Fig. S9). The most intense selective pressure was localized to nitrogen respiration gene which BLASTn/x alignment significantly annotate to the *Bacillus cereus* group (including *B. thuringiensis*). Fixed mutations (AF=1.0) were identified in the respiratory nitrate reductase alpha chain and associated subunits. This indicates a high-velocity optimization of the denitrification pathway in response to nitrogen saturation. GNAT family N-acetyltransferases and RNA methyltransferases annotated to *Bacillus* and *Klebsiella* homologs, posits that stress is re-equipping the epigenetic and translational control of the population. High-impact hits in helicase-related proteins and ParA/RecA families further support a model where the community is actively shuffling its genetic material to find viable adaptive configurations. The genomic shock also extended beyond the dominant *Bacillus* taxa into the rarer and uncharacterized members of the rhizosphere. These sample-specific consensus variants on metagenome assembled genomes (MAGs) mapped to *Nitrososphaeraceae* (Archaea) *and Candidatus Nitrosocosmicus*, indicate that the structural shifts in the ammonia-oxidizing machinery of the soil (Fig 3c). The identification of fixed hits in ATP-dependent zinc metalloproteases (FtsH) on (*Acidobacteriota*) and Type IX secretion systems mapped to *Adhaeribacter aerolatus* proves that core protein quality control and root-colonization mechanisms are being modified across multiple phyla.

The statistical benchmarking of mutational pressure against the global background (Drift Density = 0.967 mut/kb) revealed an extreme, localized genomic shock in intergenic region (promoter/regulatory space) within the inorganic treatment (Table 3). High-precision regulatory hits were not merely more frequent than stochastic mutations; they exhibited enrichment ratios several orders of magnitude higher than the genomic mean. Notably, regulatory regions for hypothetical proteins showed a staggering 2,264-fold enrichment over the background drift, while the IS200/IS605 Transposase and NAD(P)H-quinone oxidoreductase exhibited 98-fold and 77-fold enrichments, respectively. This non-random clustering within the 0–200 bp transcriptional initiation sites (Fig. 3b) confirm a targeted evolutionary strategy prioritizing rapid regulatory reprogramming of the ‘Nitrogen Stress Toolkit’ over stochastic structural protein evolution. Furthermore, we identified a significant selective sweep localized to the promoter regions of the Amino-acid permease RocC (ER = 67.2x) and NAD(P)H-quinone oxidoreductase (ER = 77.5x), (Table 3). Rather than general nutrient uptake, the targeted regulation of RocC, a specialized transporter for arginine and ornithine suggests an adaptive shift toward scavenging specific nitrogenous compounds to maintain cellular nitrogen/carbon ratios under the osmotic stress of inorganic fertilization. This is coupled with the rapid stabilization of cellular redox potential to mitigate NPK-induced oxidative stress. This scavenging strategy is integrated with the ATP-dependent zinc metalloprotease FtsH (ER = 41.3x) and the Major Biofilm Matrix (ER = 67.2x).

**Table 3.** Localized Enrichment of Fixed Regulatory Mutations under Inorganic Nitrogen Stress.

| Associated Gene (Closest Neighbor) | # Regulatory Mutation (0–200 bp Upstream) | Local Density (mut/kb) | Enrichment ratio (ER) |
| --- | --- | --- | --- |
| Hypothetical protein | 438 | 2190 | 2264.74 |
| IS200/IS605 family transposase ISBth17 | 19 | 95 | 98.24 |
| NAD(P)H-quinone oxidoreductase subunit J chloroplastic | 15 | 75 | 77.56 |
| Major biofilm matrix component | 13 | 65 | 67.22 |
| Amino-acid permease RocC | 13 | 65 | 67.22 |
| Glutamine--fructose-6-phosphate aminotransferase [isomerizing] | 13 | 65 | 67.22 |
| Spore coat assembly protein ExsA | 11 | 55 | 56.88 |
| Aspartate ammonia-lyase | 9 | 45 | 46.54 |
| Hydroxyacylglutathione hydrolase | 8 | 40 | 41.37 |
| ATP-dependent zinc metalloprotease FtsH | 8 | 40 | 41.37 |
| Global background drift density = 0.9670 mut/kb.<br>Enrichment Ratio calculated as (Local Density / Background Density). |  |  |  |

To determine the impact of mutation burden by different fertilizer treatments, we performed functional enrichment analysis on the mutation hotspots (Fig. 3d). The analysis revealed significant enrichment of mobile genetic element (MGE) mobilization and signaling regulation in the organic fertilizer treatment. Whereas inorganic treatment revealed enrichment in multidrug resistance (MDR), oxidative stress, SOS repair mutagenesis, aminoglycoside and nitrogen metabolism. These results represents that organic treatment is focused on the signaling and metabolic adaptations, whereas the inorganic treatment is focused on the structural and stress-defense mechanisms.

The MFIFM cohort analysis also revealed similar pattern on mutation burden in the genetic and intergenic regions, genomic shock, regulatory-selection signals, and gene recovery in stress and metabolic pathways during different inorganic fertilizer regimens (Fig. S10). Interestingly, we identified that bulk soil cultivator no till intensive inorganic fertilizer (BS-CT-Int) stands out as the strongest overall genomic shock treatment with highest coding mutation burden, the highest intergenic burden, and the highest number of fully recovered genes (Fig. 3e, Fig. S10a-c). At the same time, most regulatory hotspot-associated genes across all treatments still have dN/dS<1, which suggests recurrent hits are common but are not usually accompanied by strong protein-level positive selection. The functional enrichment analysis of treatments differs in total mutation burden, but the burden is repeatedly concentrated in nitrogen/phosphorus metabolism and SOS-repair-related annotations, with additional recurring signal in aminoglycoside resistance, efflux, and mobile-element associated genes (Fig. S10d). Taken together, these results suggest the dominant evolutionary response is likely regulatory rewiring and promoter/intergenic remodeling, with some parallel coding changes, rather than widespread positive selection on protein sequences.

### Inorganic fertilization expands highly conserved mobile loci a massive cryptic plasmid reservoir

To determine the genomic context of the observed high-velocity adaptation, we cross-referenced mutation hotspots with geNomad classifications. The quantitative assessment of mutation densities demonstrated a stark compartmentalization of evolutionary pressure while the core chromosome remained relatively stable, Mobile Genetic Elements (MGEs) functioned as hyper-variable hotspots (Fig. 4a). Functional profiling confirmed that these mobile hotspots carry critical survival machinery. Most notably, we identified targeted mutations in phosphonate metabolism (a thiolase involved in alternative phosphorus scavenging) hosted on the plasmid contig NODE_1029, and nitrogen assimilation (aspartate ammonia-lyase) hosted on the viral scaffold NODE_10. The presence of seven fixed mutations in the latter suggests that these Auxiliary Metabolic Genes (AMGs) are undergoing rapid diversifying selection to optimize nitrogen processing under inorganic NPK flux (25). Furthermore, we observed multiple mutations in structural stability proteins, including chromosome partitioning proteins (Smc) and IS200/IS605 family transposases (NODE_114), signifying that the mobility and maintenance of these elements are themselves under active selection (Table SX).

**Figure 4.**
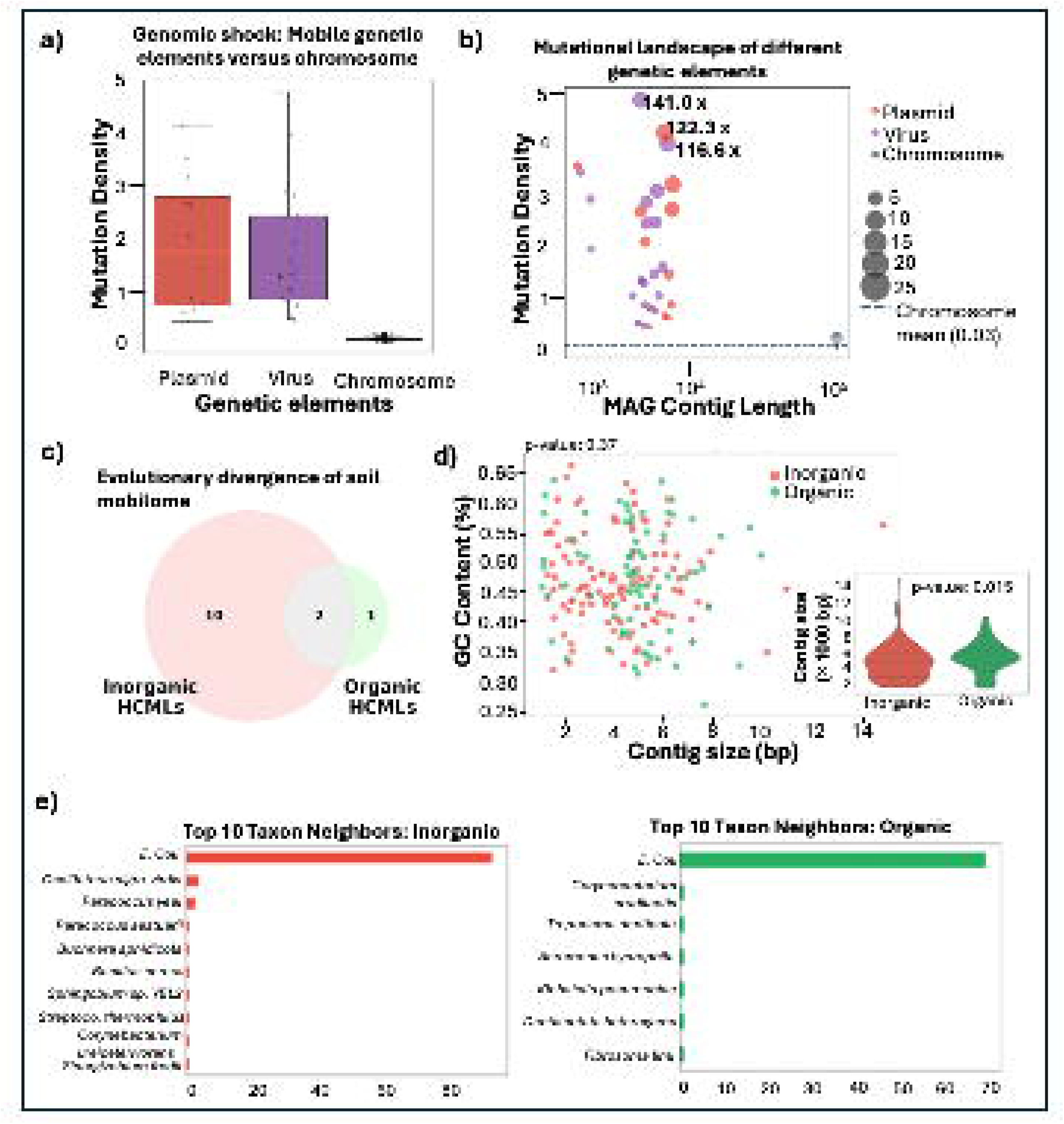
Compartmentalization of Evolutionary Velocity and the "Mobile Fortress" Effect. **(a).** Boxplot comparing the mutational load of chromosomal contigs against mobile genetic elements (MGEs). Chromosomal scaffolds exhibit high evolutionary stasis (mean = 0.034 hits/kb), while plasmids and viruses undergo a significant genomic shock, characterized by a marked increase in diversifying selection. **(b).** The mutational landscape as a function of contig length and total mutation count (bubble size). The dashed horizontal line represents the chromosomal baseline (0.034 hits/kb), highlighting the accelerated evolutionary velocity of the mobile reservoir. Primary evolutionary pioneers, including NODE_309 (Virus) and NODE_61 (Plasmid), exhibit up to a 141-fold increase in mutation density relative to the chromosomal mean. **(c).** Shared HCMLs in inorganic and organic soil mobilome. **(d).** GC content distribution of HCMLs in inorganic and organic fertilizer treatment with smaller contig size in inorganic sample (*p<0.015*). (e). The nearest neighbor taxonomic assignment of mobilome features in inorganic and organic fertilizer treatment.

Next, we explore the size of plasmid and viral genes with high mutation density. Several MAGs from inorganic fertilization sample exhibiting a state of genomic stress, with mutation densities exceeding 4.0 hits/kb representing a nearly 10-fold increase over the chromosomal background (Fig. 4b). Specifically, mobile contig NODE_309 (identified as a Viral element) and plasmid-derived contig NODE_61 emerged as the primary evolutionary pioneers, accumulating 22 and 28 mutations, respectively. Likewise, plasmid and viral gene assembled on shorter contig lengths were enriched in MFIFM cohort of inorganic fertilization (Fig. S11a-c, Fig. S12). These finding highlights that genomic shock is concentrated disproportionately on mobile genetic elements rather than chromosomes. The strongest mutation load is carried by plasmid and virus-associated nodes, especially short ones, while chromosomes provide the low-density background. That supports an interpretation where mobile elements are major hotspots of mutational turnover and possibly rapid adaptation or instability under the treatment conditions.

To identify the essential, conserved, non-redundant genetic elements which are part of coding sequence and have intense purifying selection we utilized an integrative approach. Following stringent filtering to remove co-assembly mapping artifacts, we identified a total of 185 Highly Conserved Mobile Loci (HCML) (Table SX). Fertilization treatment-unique HCMLs which drives the conservation of a distinct mobile shield (10 uniquely annotated functional families) heavily focused on stress mitigation, DNA repair, and secondary metabolism (Fig. 4c). To identify the physical architecture and evolutionary shifts of HCML, we explored the contig length and coding content. HCMLs were largely sequestered on minimalist genetic vehicles, functioning as cryptic circular elements that lack canonical conjugation machinery but maintain high stability within the community (25) (Fig. 4d). This structural divergence occurred despite a lack of significant difference in overall host taxonomic signature (GC content) and predicted mobility potential. The nearly identical GC% profiles across treatments but smaller inorganic contigs suggest that while the genetic vehicles adapt in size and complexity to accommodate different functional cargo such as the larger defense suites seen in the inorganic samples, they are maintained within a similar taxonomic host framework (Fig 4d).

To identify the biological sources of genetic vehicles, we performed MASH-based nearest-neighbor identification on the extracted contigs (26). While both treatments were heavily populated by genetic elements with high similarity to Copiotroph generalist species like *Escherichia coli*, the remaining taxonomic profiles exhibited distinct treatment-driven divergence (Fig. 4e). The inorganic mobilome showed enrichment in Oligotrophs neighbors associated with chemical versatility, such as *Paracoccus* and *Oscillatoria*, further supporting a specialized mobilome geared toward inorganic chemical adaptation. In contrast, the organic mobilome was associated with a broader suite of Copiotrophs environmental bacteria, including *Fibrisoma limi* and *Aeromonas hydrophila*, reflecting the higher ecological heterogeneity typical of organic-amended soils (Fig. 4e). This taxonomic shift suggests that organic fertilization supports a more diverse, generalist mobilome, whereas inorganic fertilization selects for a specialized, structurally distinct, and taxonomically focused set of conserved mobile elements capable of navigating extreme agricultural stress

### Implication of inorganic fertilization on plant health regulatory response

To explore the maize root genes impacted by the inorganic fertilizer treatment, we performed RNA-Seq data analysis in the publicly available dataset (27). The transcriptome profiling revealed a strong treatment effect in the Inorganic treated roots. Principal component analysis clearly separated control, inorganic, and mixed samples, with PC1 and PC2 explaining 48.08% and 24.63% of the variance, respectively, indicating distinct and reproducible expression states across treatments (Fig. 5a). Differential expression analysis identified 580 upregulated and 1,385 downregulated genes in the inorganic condition (*log2FC > |2|, adj-Pval < 0.05*), consistent with the asymmetry observed in the volcano plot and heatmap clustering (Fig. 5b). Functional enrichment analysis showed that inorganic fertilizer treated upregulated genes were predominantly associated with carbohydrate metabolism, chitin catabolism, and cell wall macromolecule catabolic processes, whereas downregulated genes were enriched for oxidative stress response, hydrogen peroxide catabolism, peroxidase activity, heme binding, and plant-type secondary cell wall biogenesis (28) (Fig. 5c, Fig. S13). Additional enriched terms, including DNA-binding transcription factor activity, glutathione transferase activity, glutathione metabolic process, monooxygenase activity, and oxidoreductase activity, further support activation of detoxification and stress-associated metabolic pathways. Consistent with this pattern, the updated DEG annotation identified 64 differentially expressed regulators in total, including 38 upregulated and 26 downregulated regulators (Table SX). The regulator-pathway interaction network showed that transcription-centered pathways were the dominant regulatory hubs, with regulation of DNA-templated transcription and DNA-binding transcription factor activity containing the largest regulator overlap and being driven entirely by upregulated regulators (DREBs, NAC61, NAC90, WRKY, and MYB59 like), whereas iron ion binding, heme binding, monooxygenase activity, and related oxidoreductase pathways were associated mainly with downregulated regulators (AP2/ERF family) (Fig 5c). Such regulatory rebalancing is consistent with the downstream enrichment of oxidative stress response, hydrogen peroxide catabolism, peroxidase activity, heme binding, and cell wall-related processes (29, 30). These results support the interpretation that maize roots under inorganic fertilization undergo coordinated transcriptional reprogramming to integrate nutrient availability with redox homeostasis, structural adjustment, and stress adaptation.

**Figure 5.**
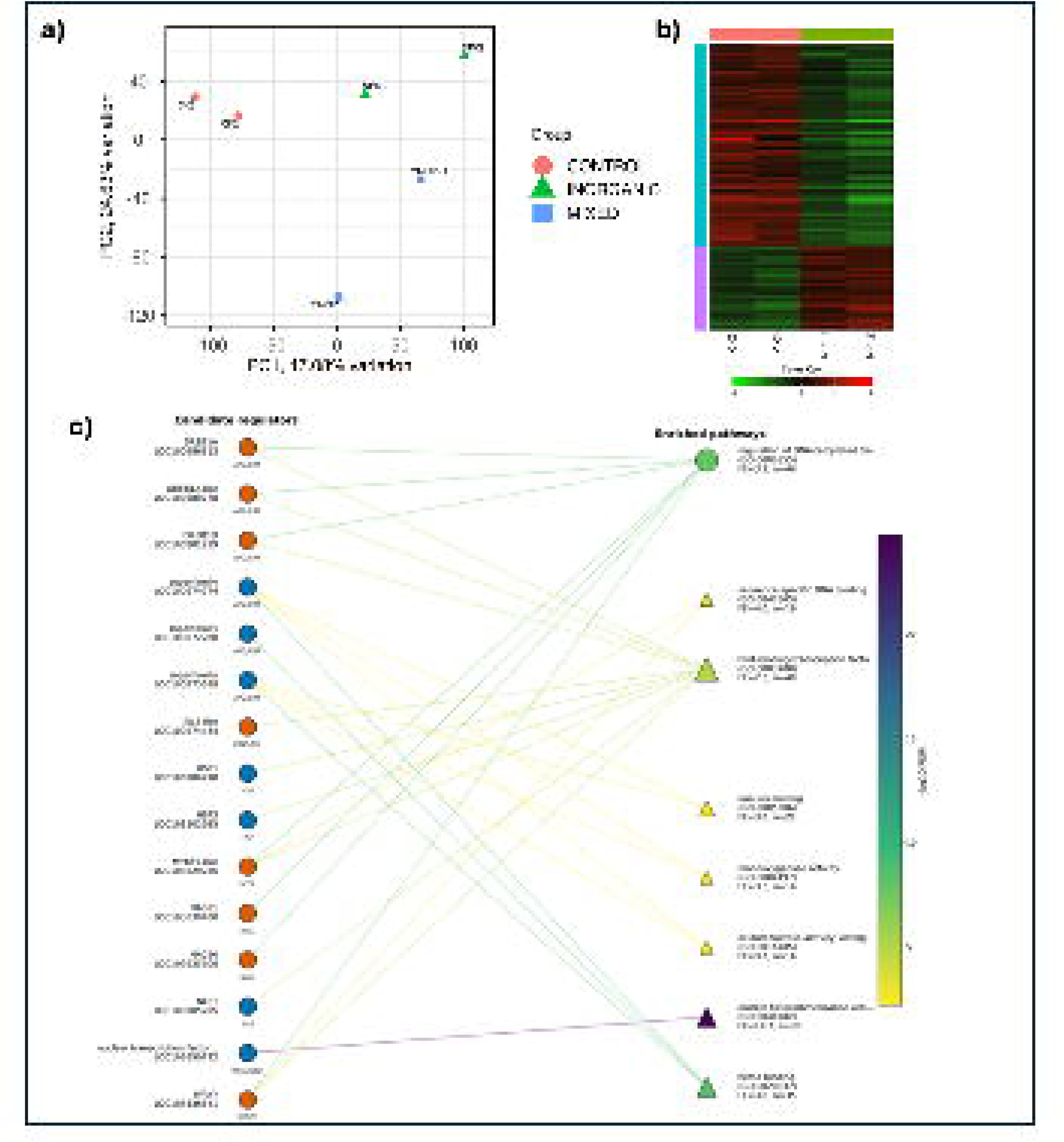
Impact of inorganic fertilization on maize root. **(a).** The principal component analysis (PCA) of root RNA-Seq data demonstrate a clear separation among control and inorganic samples. **(b).** The heatmap of differentially expressed genes (DEGs) a total of 580 upregulated and 1,385 downregulated genes in the inorganic condition (*log2FC > |2|, adj-Pval < 0.05*) with red indicating relatively higher expression and green indicating relatively lower expression. (c). Network plot of enriched pathways and the significantly expressed regulators associated with those pathways during inorganic fertilizer treatment on maize root. Regulator nodes are colored by direction of differential expression, and pathway nodes encode ontology, gene-set size, and enrichment significance.

### Shared Stress Response During Inorganic Fertilization on Microbes and Plant

Pangenomic analysis reveals that fertilization regimes fundamentally dictate the metabolic costs and survival strategies of the maize rhizosphere microbiome, driving a 1.9-fold expansion of the accessory genome under inorganic (NPK) fertilization compared to organic management (Table 4). This genomic shift under the inorganic regime is characterized by a 98.24-fold enrichment in IS200/IS605 family transposases taxonomically anchored to dominant *Bacillus cereus* rhizosphere pioneers, selecting for a survivalist community that prioritizes structural defense such as cell-wall integrity (*WalR*) and antimicrobial mechanisms (*YxdM/L*) to mitigate osmotic stress and chemical volatility. Conversely, organic fertilization promotes a synergistic mutualism focused on sustainable nutrient mobilization, enriching pathways for organic nitrogen recycling (*DppE*), iron scavenging (*FeuC*/*YfiY*), and rapid root colonization via spore germination (*YndE*). Remarkably, despite these divergent adaptive trajectories, both regimes tightly maintain a highly conserved core plant growth-promoting potential, marked by a massive enrichment in isonitrile hydratase (*LFC > 6.7*) for phytohormone biosynthesis, demonstrating that direct hormonal stimulation of root architecture remains a resilient functional baseline irrespective of abiotic agricultural stress.

**Table 4.** Functional annotation of maize rhozbium microbiome pangenome differences in organic and inorganic fertilization.

| Functional Category | Representative Gene/Pathway | Primary Group | Benefit to Maize / Rhizosphere Role |
| --- | --- | --- | --- |
| Hormonal Stimulation (IAA) | Isonitrile hydratase | Both | Potential synthesis of Indole-3-acetic acid (IAA). Directly promotes root elongation and lateral root branching in maize. |
| Nitrogen Scavenging | DppE (Dipeptide-binding) | Organic | Recycles organic nitrogen from root exudates and fertilizer. Increases N-availability in the immediate root zone. |
| Iron Mobilization | FeuC / YfiY (Siderophores) | Organic | Sequesters iron for the plant while inhibiting soil-borne pathogens. Crucial for maize chlorophyll and photosynthesis. |
| Osmotic Tolerance | WalR / Peptidoglycan deacetylase | Inorganic | Protects the bacterial cell from high salt concentrations (NPK salts). Ensures survival of beneficial microbes in high-input soil. |
| Persistence & Resilience | Spore germination proteins (YndE) | Organic | Allows bacteria to survive nutrient fluctuations and rapidly colonize roots as soon as maize seeds germinate. |
| Chemical Defense | YxdM / YxdL (ABC Transporters) | Inorganic | Provides resistance against antimicrobial peptides. Helps the community maintain its niche in a highly competitive rhizosphere. |

To identify biological processes jointly represented in the host transcriptome and microbial functional repertoire under inorganic treatment, we integrated host-transcriptome and microbial pangenome crosstalk analysis under inorganic treatment (Fig. 6). Regulation of DNA-templated transcription emerged as the largest microbial-centered module, driven by the up-regulation of five host transcription factor loci (*MYB48*) alongside a massive microbial regulatory reprogramming involving 21 genes encoding sigma factors, response regulators, and transcriptional repressors. Conversely, the response to oxidative stress was heavily host-dominated, featuring the robust down-regulation of 15 host peroxidase-related loci (e.g., *LOC100191217* and *LOC103633557*) contrasted against a narrow microbial defense investment restricted to protein repair and stress-tolerance proteins (*FtsH* and *MsrA/MsrB*) (31). Iron ion binding functioned as a balanced, intermediate response axis linking 10 host genes including down-regulated regulators (*LOC100273589* and *LOC100274274*) with 4 microbial proteins tied to cofactor biosynthesis and oxidoreductive metabolism, reflecting a coordinated shift in micronutrient handling and electron transfer. Finally, the cell-surface boundary was represented by the extracellular region pathway, a highly localized module driven predominantly by 4 microbial proteins against a single host locus. Collectively, these findings demonstrate that while inorganic chemical stress reorganizes the holobiont interactome around tightly conserved regulatory, oxidative, and metal-associated pathways, the execution of these functional themes is highly compartmentalized, with the host mounting a broad antioxidant defense while the microbiome coordinates systemic transcriptional and extracellular adaptation.

**Figure 6.**
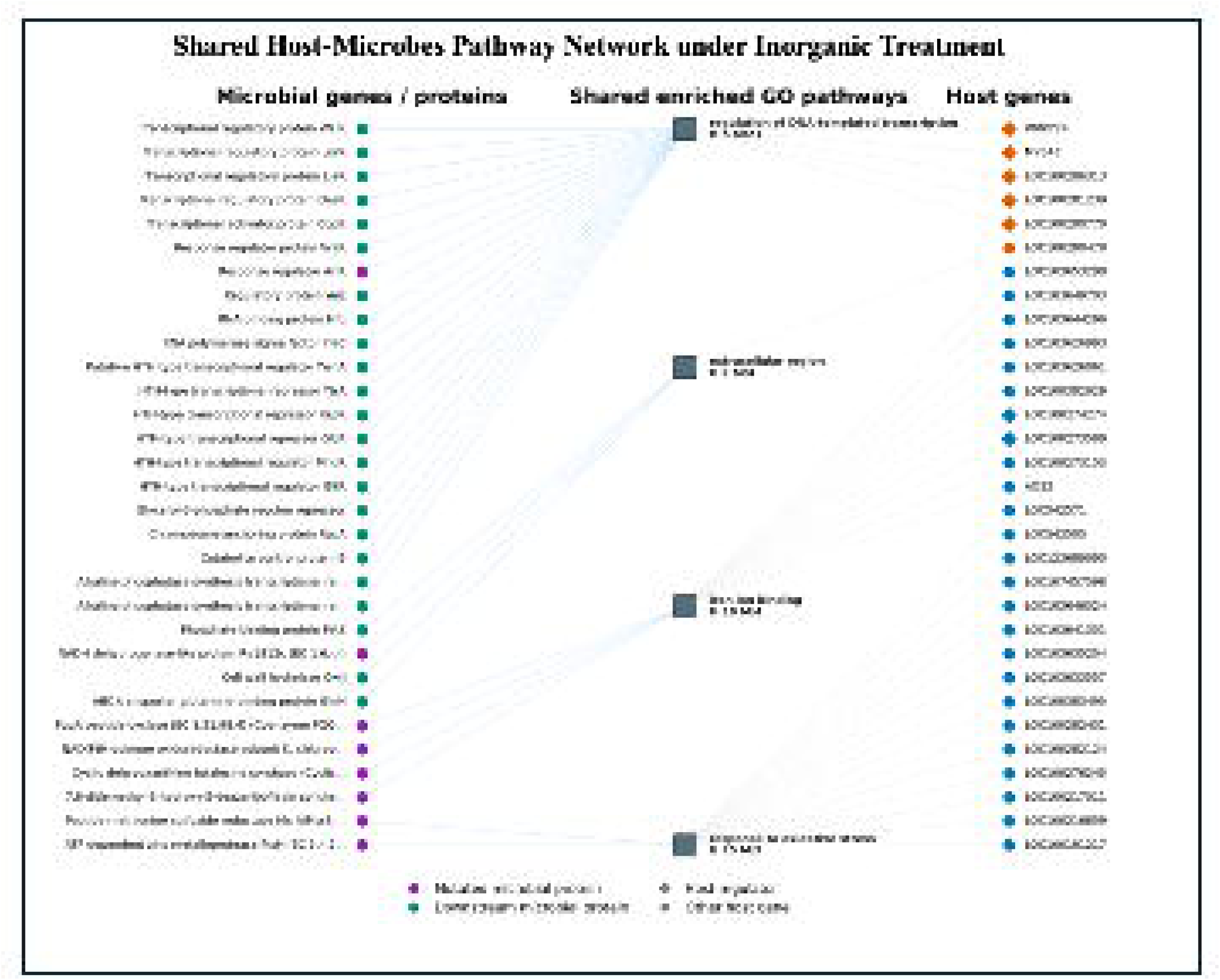
Shared host-microbe network under inorganic fertilizer treatment in maize roots. Nodes represent host genes, microbial genes/proteins, and shared enriched GO pathways, with edges indicating pathway membership. The network retained four shared pathways including regulation of DNA-templated transcription, extracellular region, iron ion binding, and response to oxidative stress. Transcription is the most microbe-enriched module, linking 21 microbial proteins with 5 upregulated host regulators, whereas oxidative stress is the largest host-centered module, linking 15 downregulated host genes with 2 microbial proteins. Iron ion binding connected 10 host genes and 4 microbial proteins, and extracellular region linked 1 host gene with 4 microbial proteins.

## Conclusion

In conclusion, our study provides a high-resolution, population-genomic paradigm shift in our understanding of how anthropogenic practices reshape the soil microbiome. By moving beyond traditional taxonomic and gene-abundance profiling, we have demonstrated that long-term soil management regimes act as powerful evolutionary forces that drive resident rhizosphere bacteria into separate, highly specialized evolutionary trajectories. Our findings resolve a fundamental evolutionary paradox within intensively managed agricultural systems. Under chronic inorganic fertilization, microbial populations do not undergo widespread, disruptive structural protein evolution despite facing severe genotoxic and osmotic stress. Instead, they deploy an elegant regulatory mechanism, strategically concentrating mutations within the immediate 0–200 bp promoter zones of critical stress-defense and metabolic genes. This genomic defense is bolstered by a structural bifurcation across genomic compartments, wherein the extra-chromosomal mobilome expands its repository of highly conserved mobile loci on cryptic circular vehicles, serving as a rapid-response functional vault to bypass systemic environmental repressions. The intensive regulatory modifications force the microbiome to transition from a dynamic, mutualistic hub of plant-microbe cooperation into an isolated defensive enclave. Ultimately, these insights emphasize that the successful deployment of next-generation, microbially mediated sustainable agriculture cannot rely solely on managing microbial abundance or composition.

## Methods

Refer to supplementary files.

## Author contributions

A.G.: Conceptualization, workflow development, documentation, scripting, analysis, visualization, manuscript writing, planning, management, and coordination of the soil metagenome research program. B.M. Workflow Automation and scalable implementation, manuscript review and writing, planning, management, and coordination of the Host transcriptome and integration research program. All authors discussed the results, critically reviewed the manuscript and provided feedback.

## Data and materials availability

All data needed to evaluate the conclusions in the paper are present in the paper and/or the Supplementary Materials. Additional data related to this paper may be requested from the authors. The *MAGmutome* workflow is available on the GitHub.

