## Supplemental Text for "Chronic Inorganic Fertilization Shifts Evolutionary Trajectories and Induces a Regulatory Shield in Maize Rhizosphere Bacteria"

**Supplemental Figures**

**Supplemental Figure legends**

**Materials and Methods**

**Supplemental Figures.**

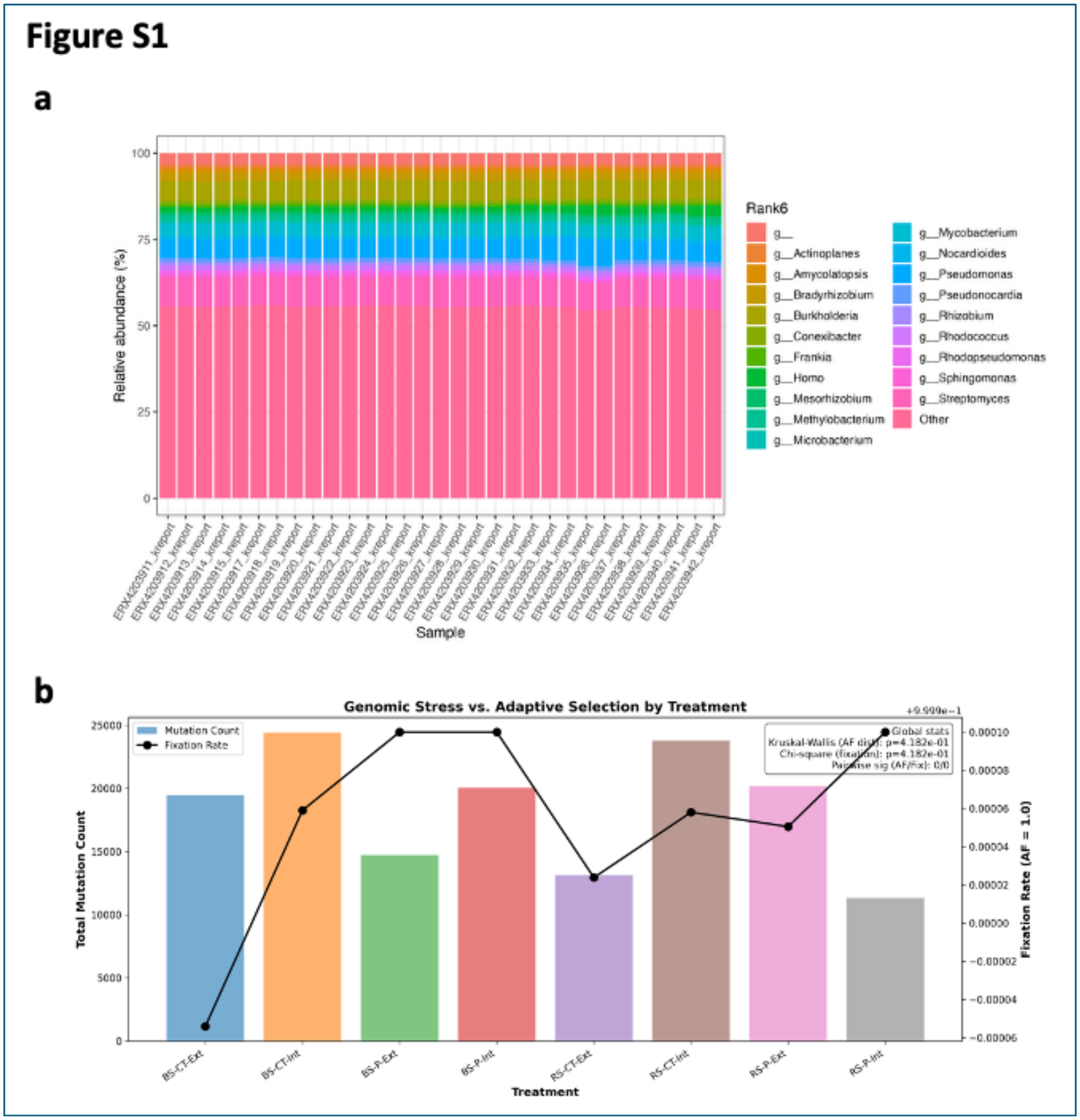


**Figure S1. The multi-factorial inorganic fertilizer management (MFIFM) cohort metagenomic data analysis. (a).** The Genus relative abundance assigned by the *MAGmutome* framework. **(b).** The total mutation count and Fixation rate in different treatment groups including bulk soil (BS) and root-affected soil (RS). Treatments comprised mould-board ploughing (P) vs. cultivator treatment (CT, reduced tillage) in combination with intensive (Int) vs. extensive (Ext) nitrogen fertilization.

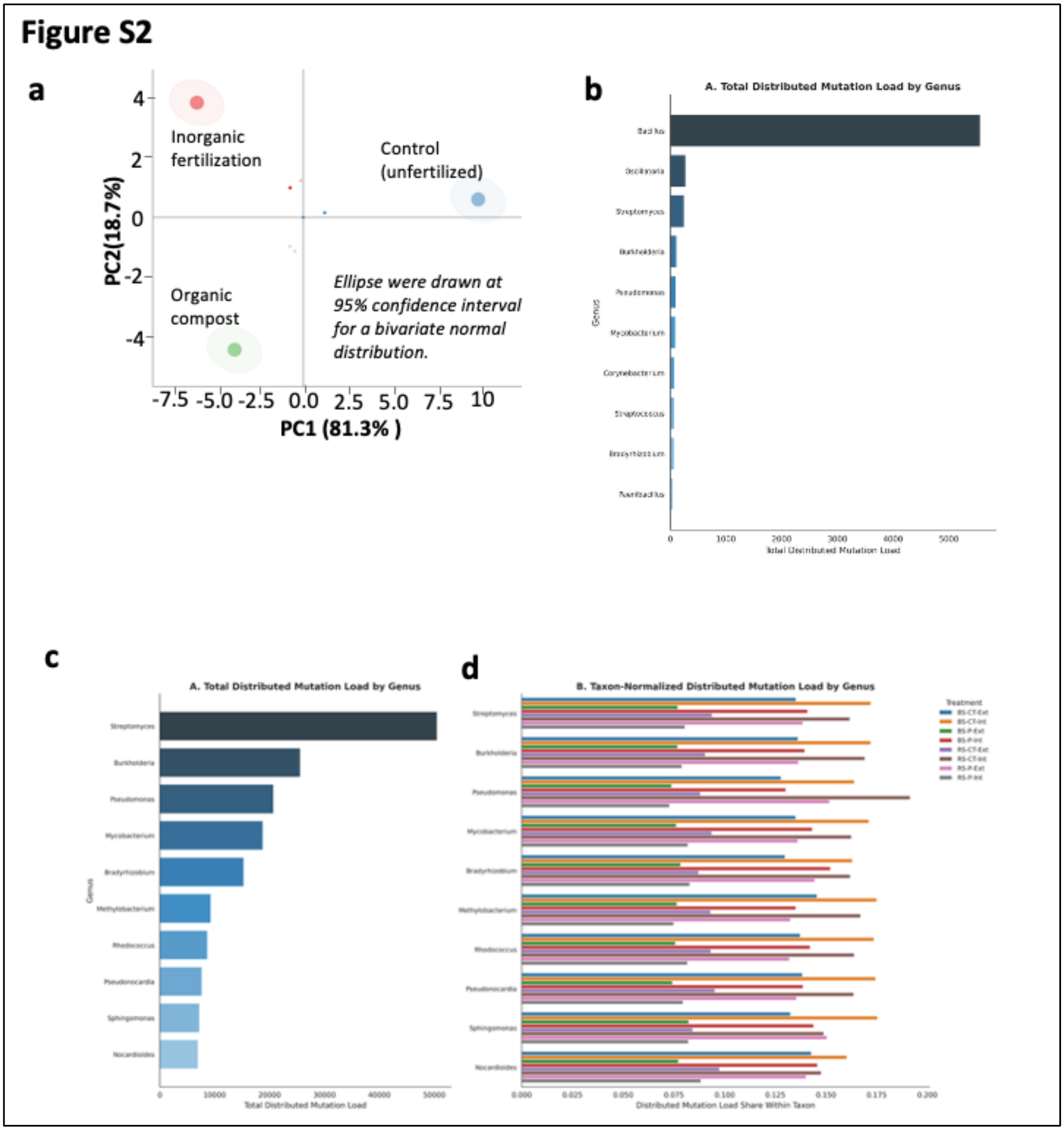


**Figure S2. Microbial adaptive survival under fertilizer treatment. (a).** The Principal Component Analysis (PCA) and ordination of bacterial evolutionary trajectories (*dN/dS* profiles) across soil management systems. Ordination plot demonstrating the distinct segregation of population-level selective regimes under Control, Inorganic NPK, and Organic compost fertilization practices. Principal Component 1 (PC1) accounts for 81.3% of the total variance, reflecting the global selection intensity across the microbial population, while Principal Component 2 (PC2) captures 18.7% of the variance, representing treatment-specific functional divergence. **(b).** Total Mutation load in top 10 genus taxon under the organic, inorganic, and control treatment. **(c).** Total Mutation load in top 10 genus taxon under the MFIFM cohort. **(d).** Total Mutation load in top 10 genus taxon under the MFIFM cohort in each treatment.

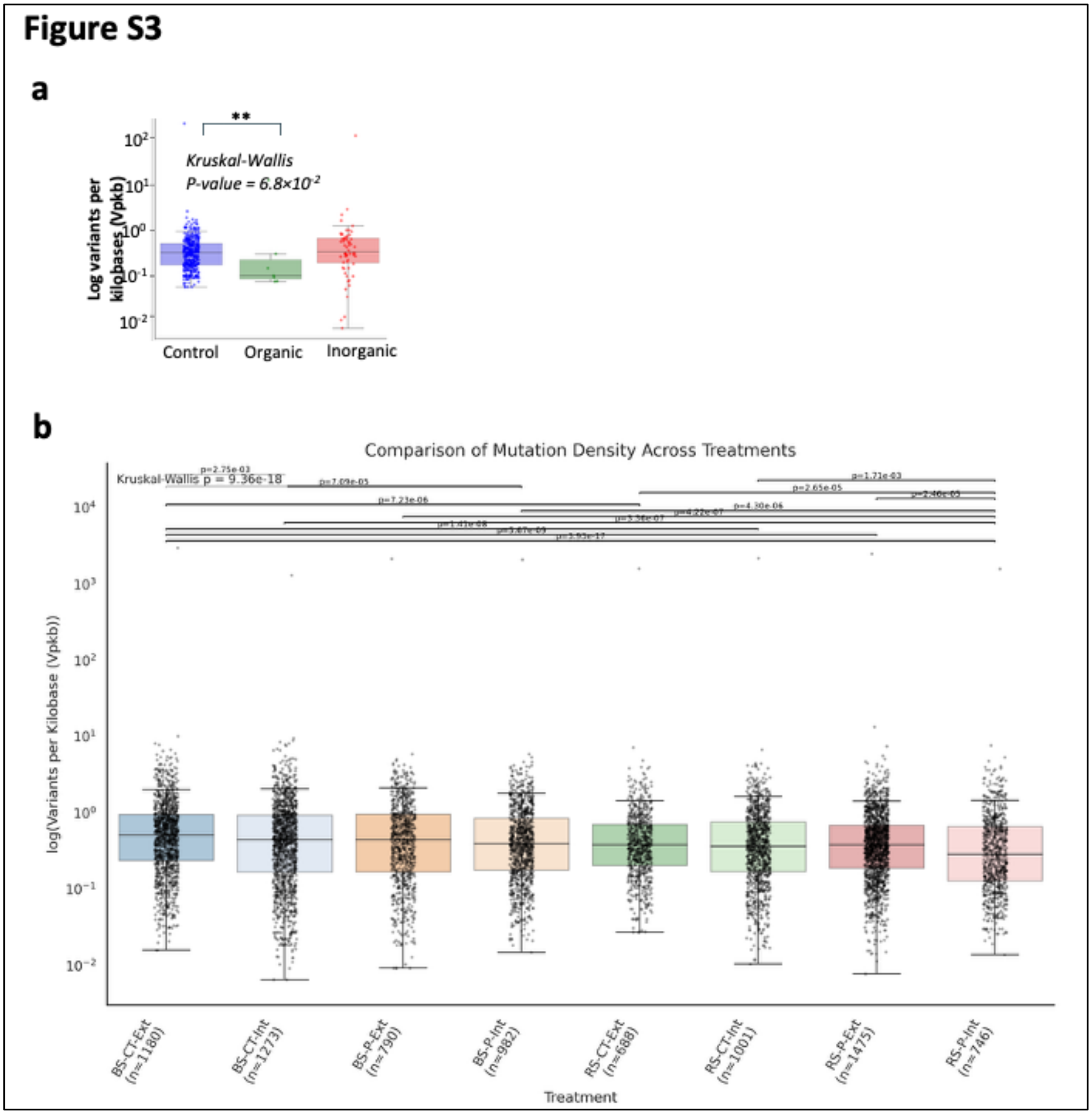


**Figure S3. The** mutational velocity (Vpkb) in the Organic, Inorganic, and Control treatment **(a)** and The MFIFM cohort treatments **(b)**. The significant differences are marked by *P-value < 0.05*.

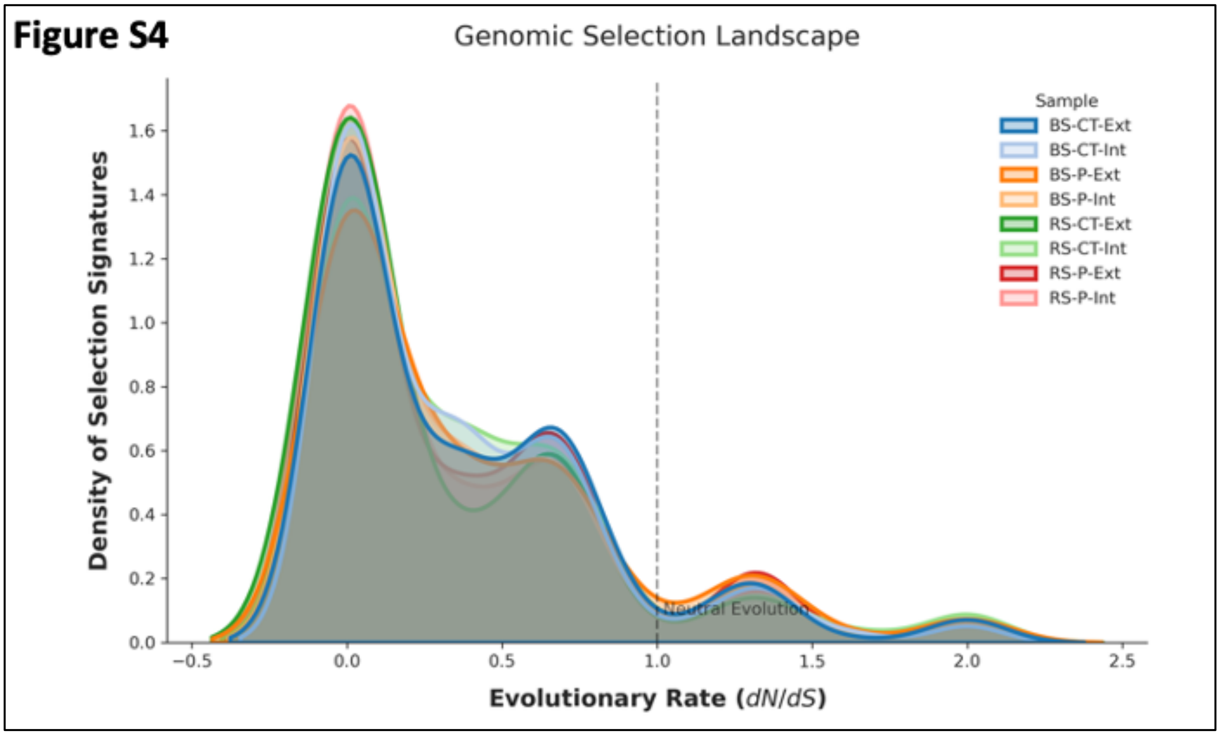


**Figure S4.** Genomic selection density landscape in different treatments of MFIFM cohort.

**Figure S5**

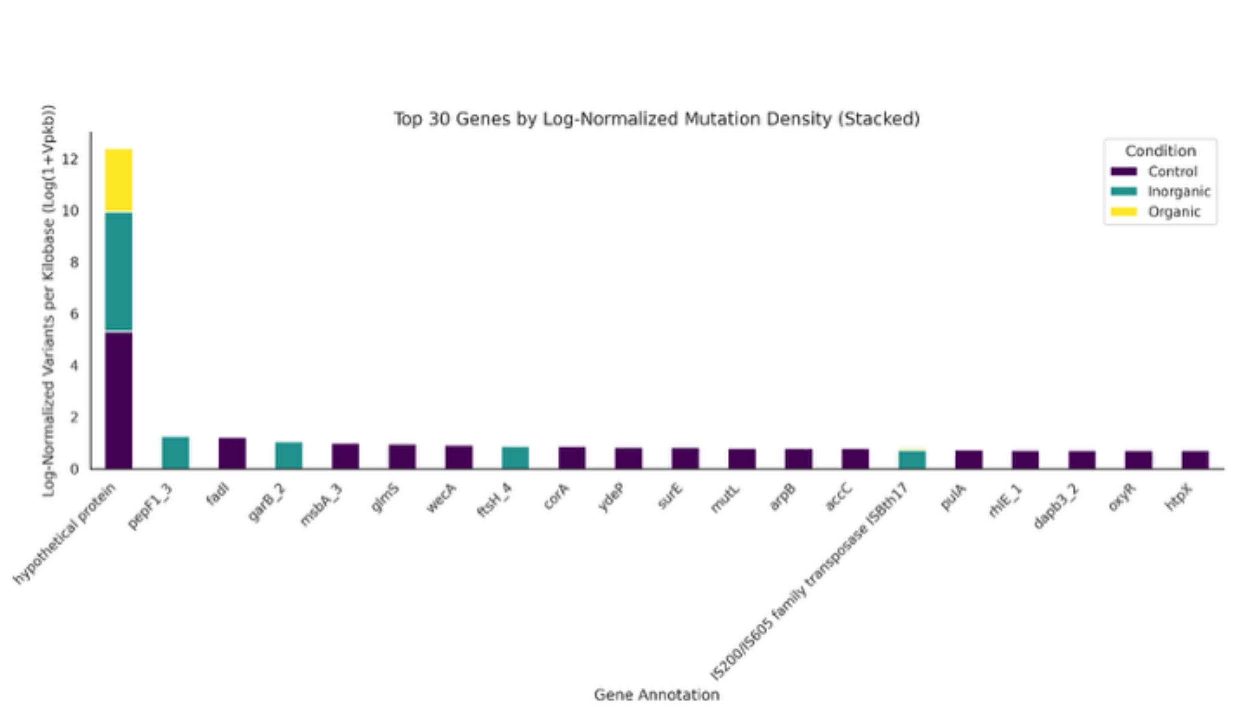


**Figure S5. Chromosomal genes have very high mutations. (a)** High volume of Mutation in genes in different fertilizer treatments including Control (dark-blue), Inorganic (light-blue), and control (yellow).

**Figure S6**

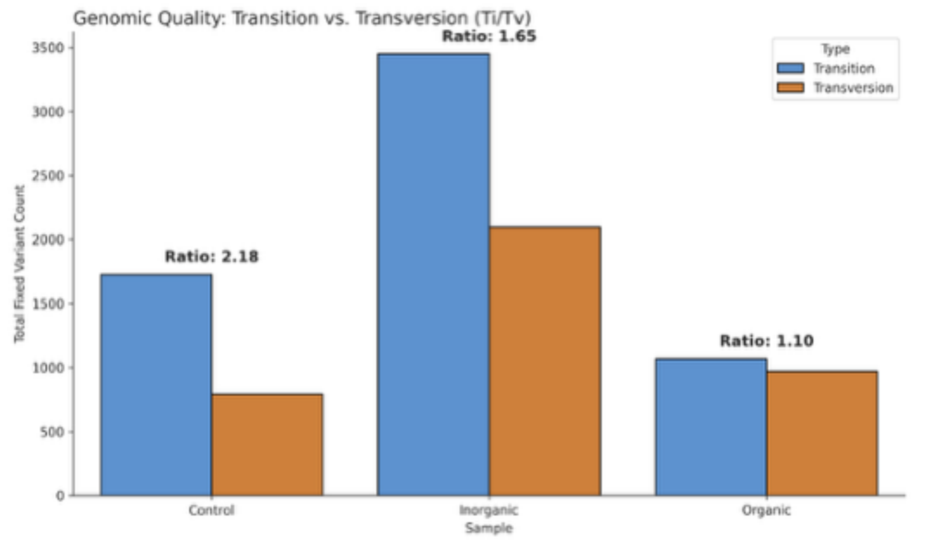


**Figure S6. The Total number of fix variants in different fertilizer treatments and the high enrichment of transitions compared to transversions in the Control and inorganic treatment.**

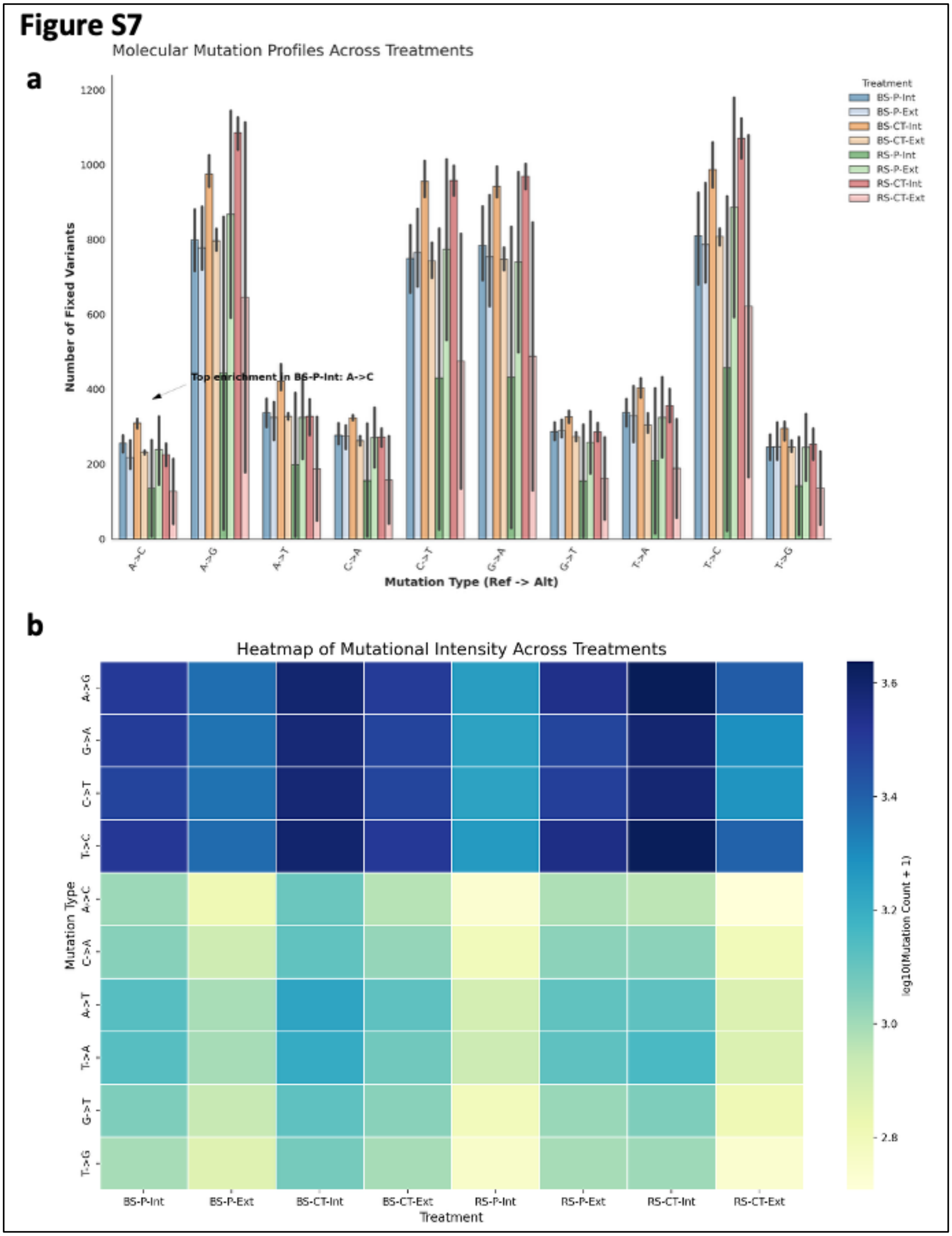


**Figure S7. Total number of fixed variants (a) in several mutation types including transition and transversion events. (b). Mutation intensities heatmap of single-base substitutions (log-transformed counts).**

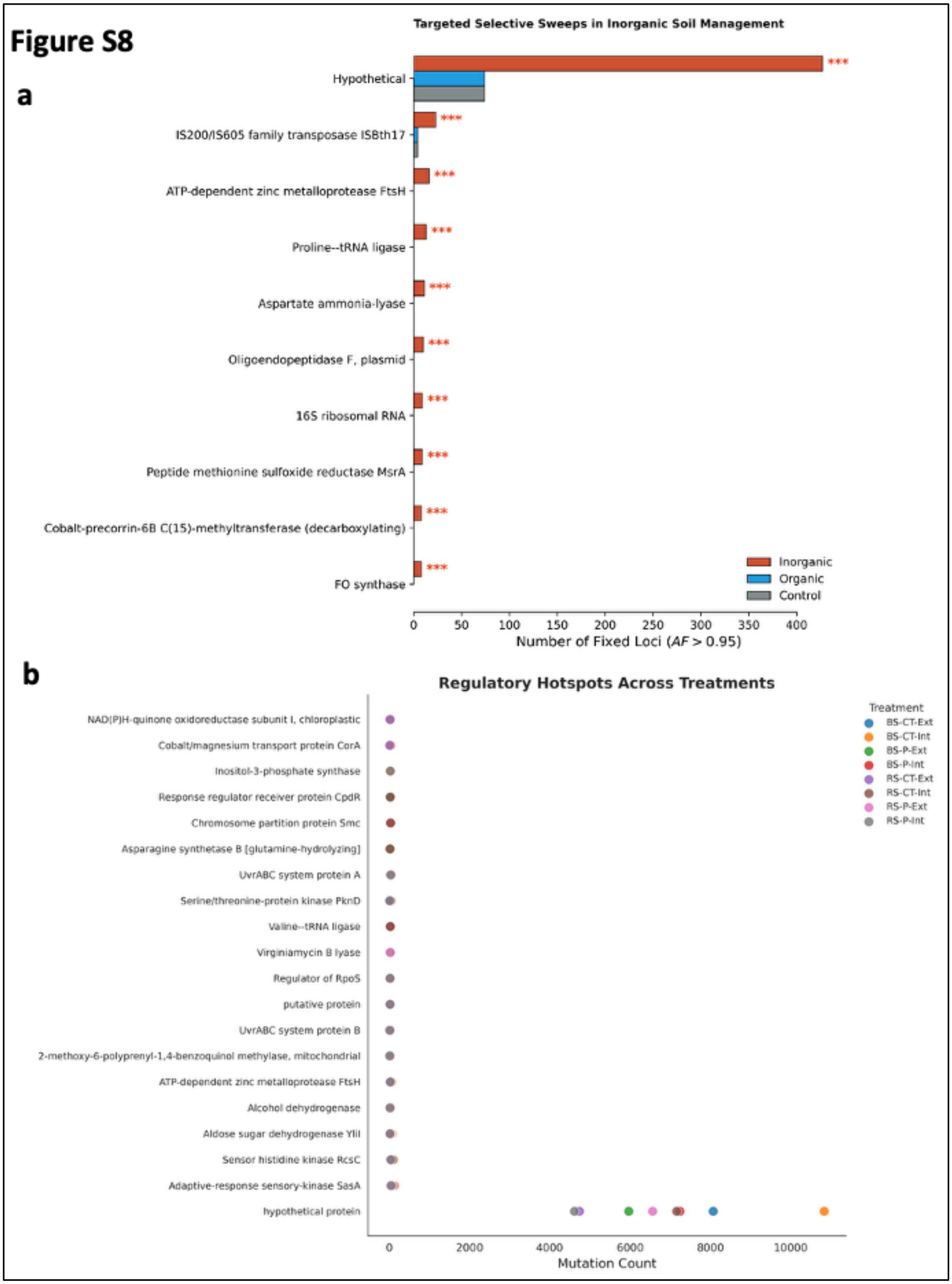


**Figure S8. Selective sweep of regulatory hotspots. (a)** Significantly high number of Fixed loci of regulatory and coding regions with allele frequency (AF) > 0.95 in inorganic treatment vs organic and control. Organic treatment exhibits a total absence of these regulatory hotspots. **(b).** Total mutated regulatory hotspot genes in MFIFM cohort treatments.

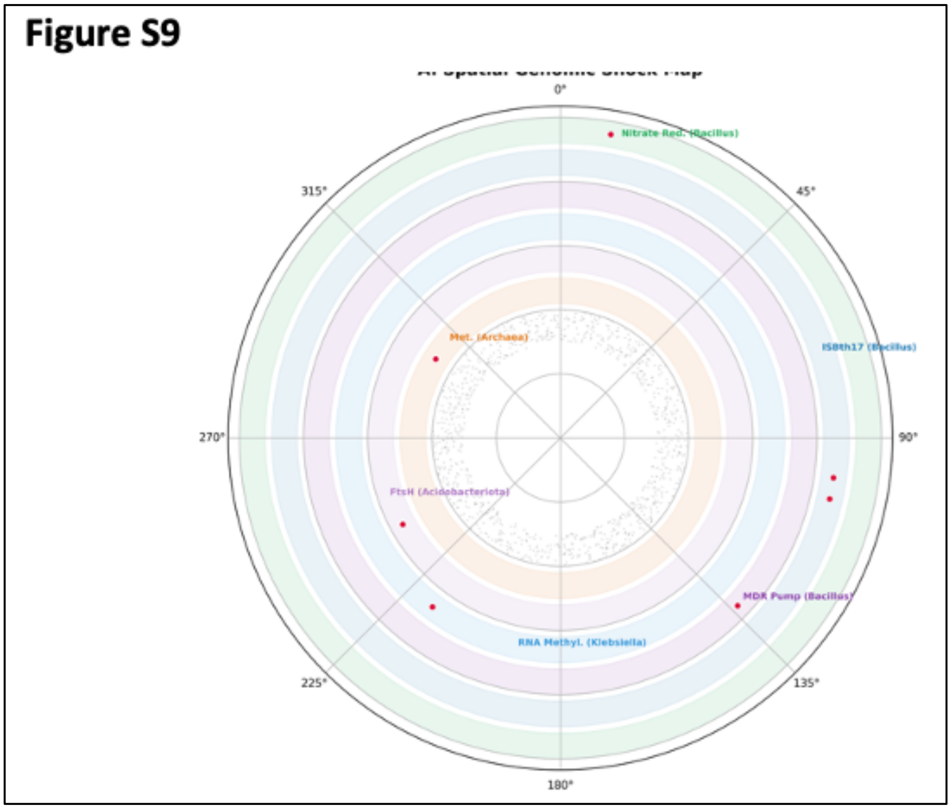


**Figure S9. Circular plot of fixed mutations annotated on significant taxon.**

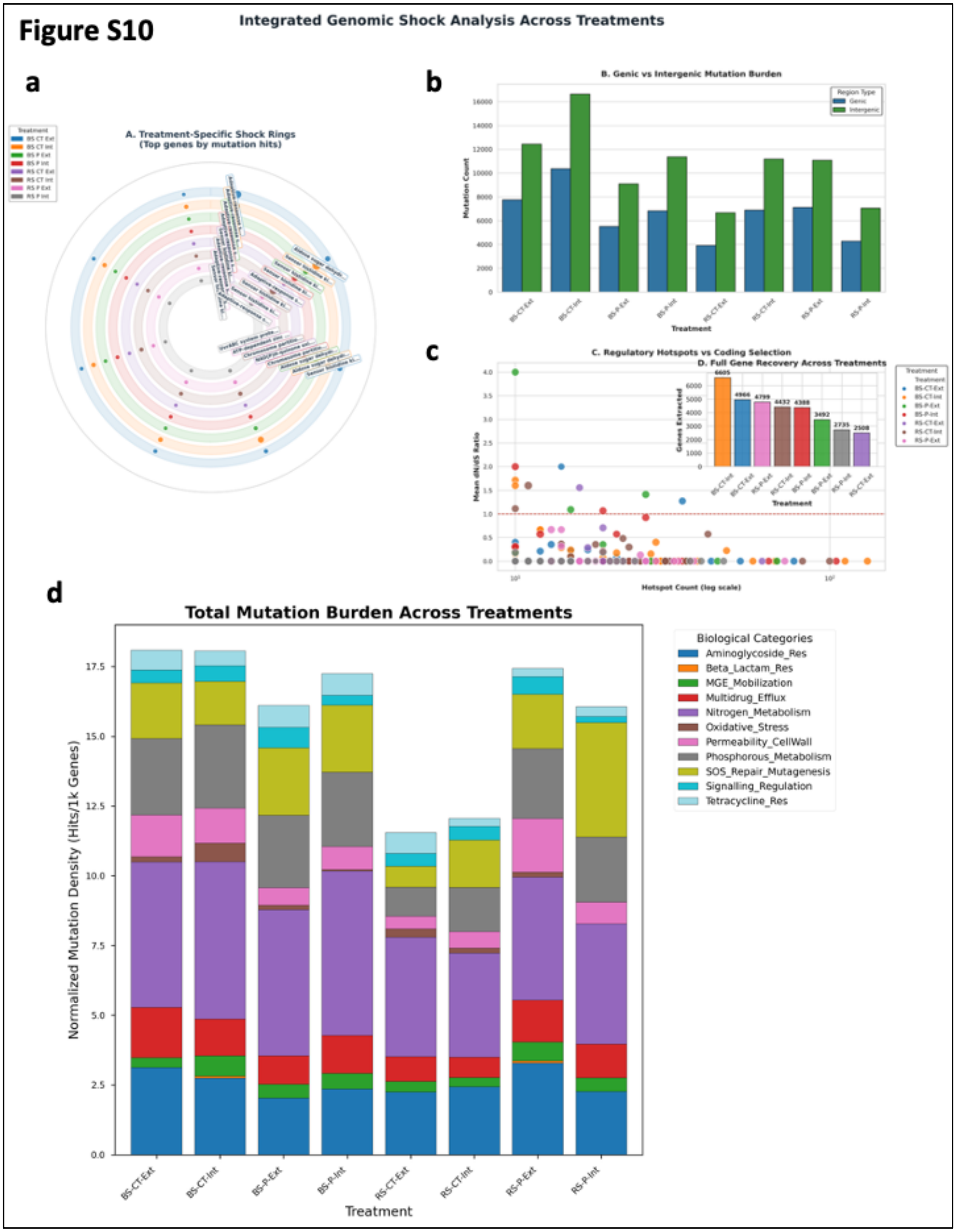


**Figure S10. (a-d) Spatial localization of fixed mutations in regulatory hotspots and biological functional categories in MFIFM cohort.**

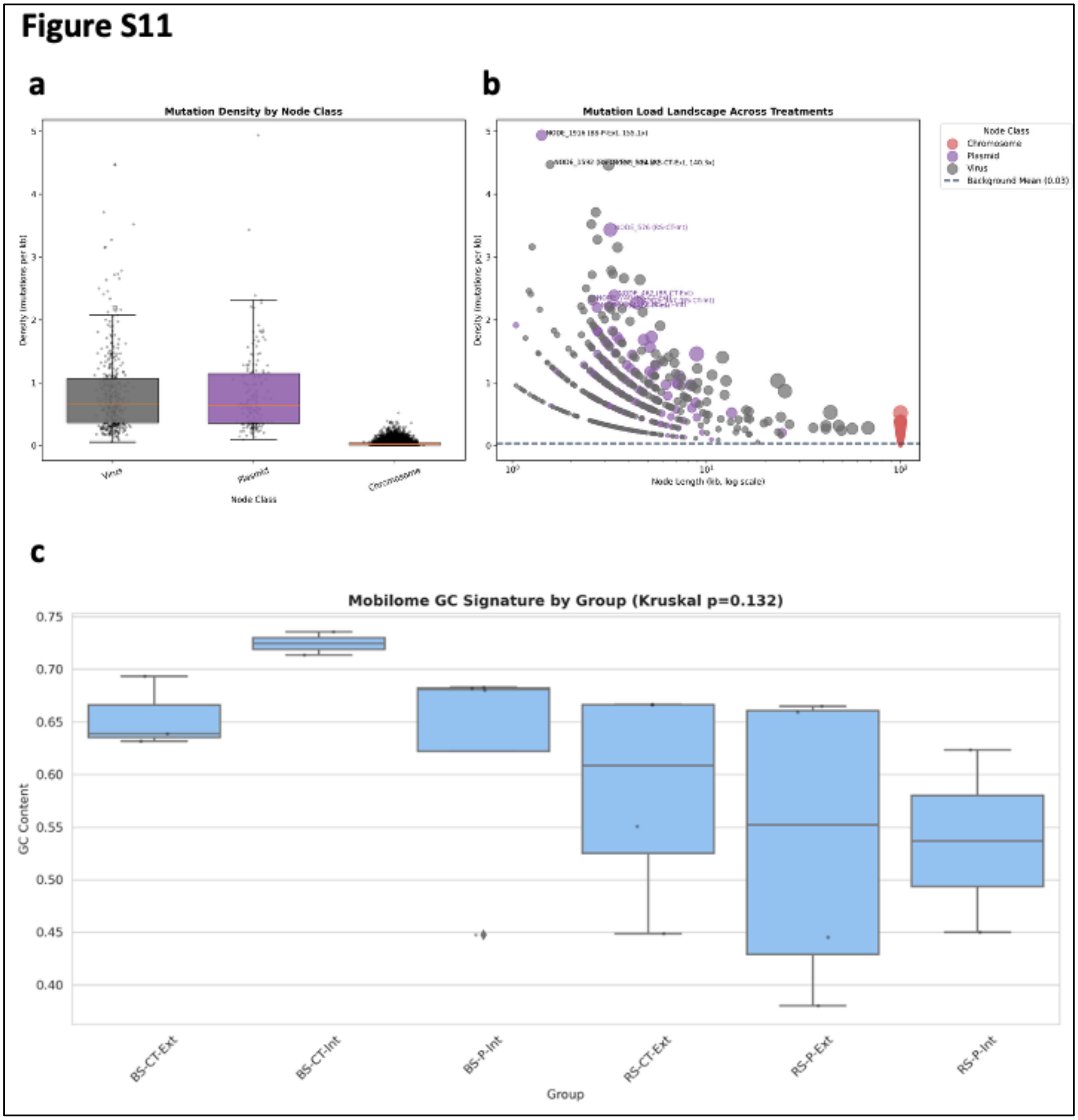


**Figure S11: Mutational load and compartmentalization in MFIFM cohort. (a).** Mutational load and compartmentalization of bacterial chromosomal and plasmids and viral genes. **(b).** The contig lengths of high mutational density in chromosomal and plasmids and viral genes. **(c).** The GC content distribution of HCMLs in MFIFM cohort treatment.

Figure S12

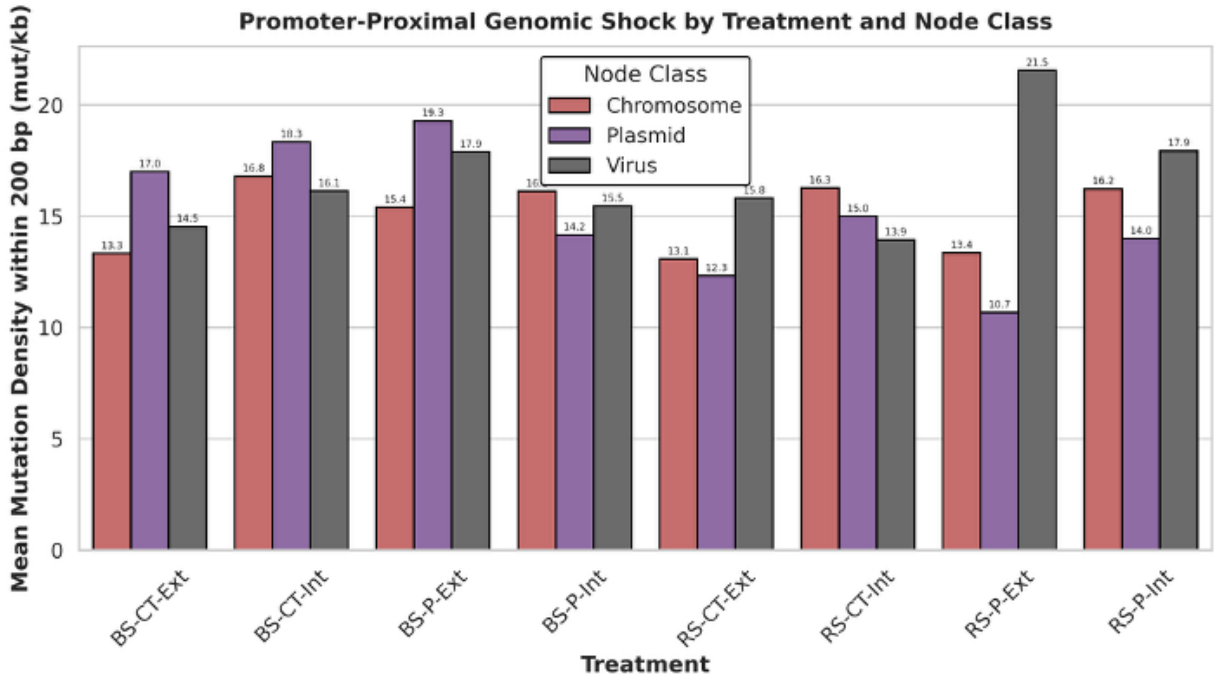


**Figure S12. The mutational density of bacterial chromosomal and plasmids, and viral genes within 200bp of promoter region.**

**Figure S13**

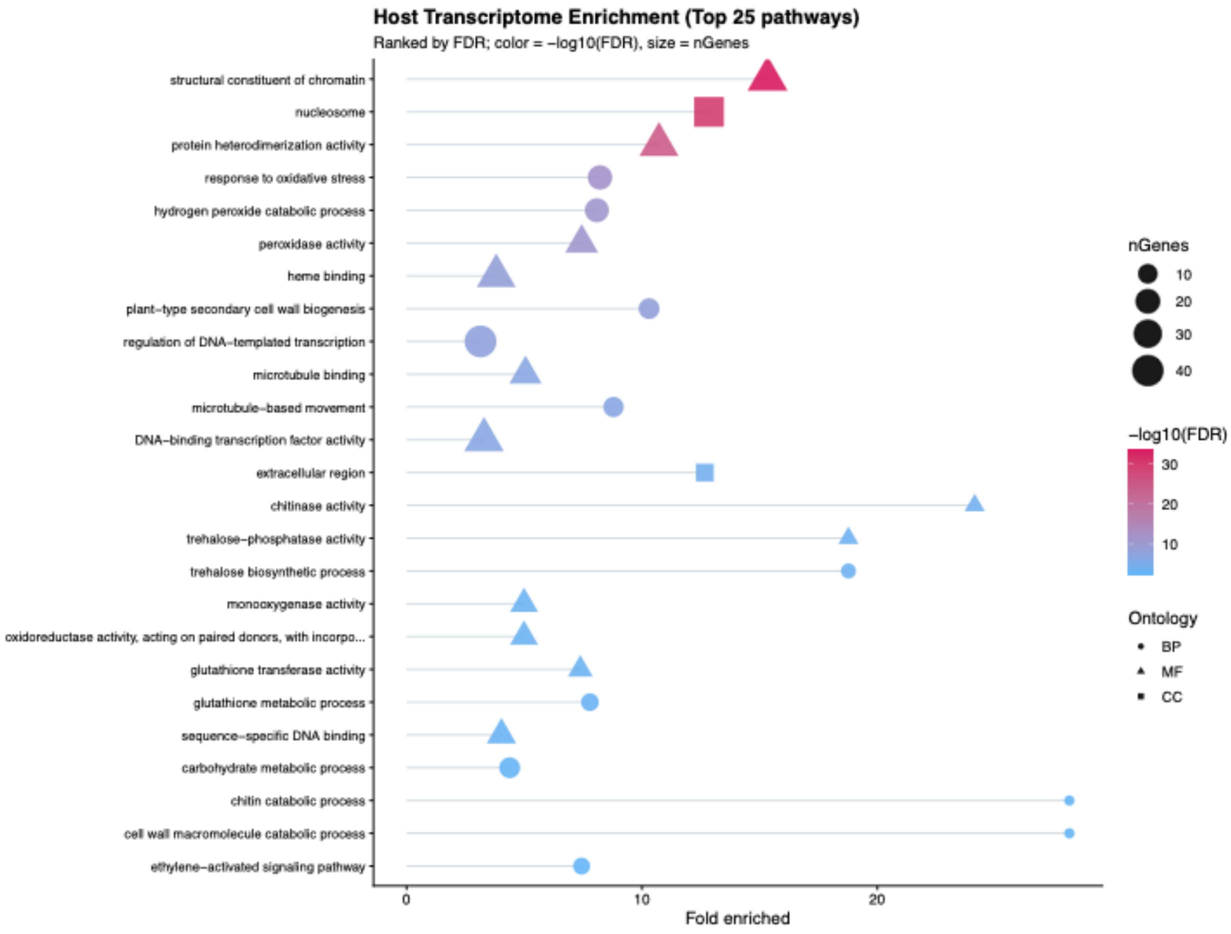


**Figure S13: The gene ontology (GO) of maize root significantly up- and down-regulated Differentially expressed genes.**

### Materials and Methods

#### Data Acquisition and metagenomic assembly

Raw shotgun metagenomic reads were obtained from the maize rhizosphere study conducted by Enebe and Babalola (1), focusing on three distinct agricultural treatments: high organic compost (CP8; 8 t/ha), high inorganic NPK fertilization (N2; 120 kg/ha), and an unfertilized control (Cn0). As established in the source study, these treatments represent a gradient of phosphorus (P) availability and soil acidification, with $N_2$ treatment previously linked to a significant repression of P-cycling gene abundance. The NCBI Sequence Read Archive (SRA) under BioProject PRJNA607213. Reads from the Inorganic (N2) and Organic (CP8) populations and Control were used in this study. Quality control was performed using FastQC and MultiQC to ensure mean base quality (Q > 30)(2, 3). Raw metagenomic reads were processed using the nf-core/mag (v4.0.0) pipeline(4, 5). To ensure high-quality assembly, reads were digitally normalized using BBNorm and quality trimmed(6). Taxonomic profiling of raw reads was performed using Centrifuge database to identify initial community composition(7). De Novo assembly was performed using SPAdes in metagenomic mode (--meta)(8). To maximize the recovery of high-quality Metagenome-Assembled Genomes (MAGs), we employed a multi-tool binning and refinement strategy: first the initial binning was performed using CONCOCT and MetaBAT2, next bins were refined using DAS Tool with a high-stringency score threshold of 0.3 (9-11). Bin completeness and contamination were assessed using CheckM and BUSCO utilizing bacteria lineage database(12, 13). For the multi-factorial inorganic fertilizer management (MFIFM) the dataset used was extracted from NCBI Bioproject ID PRJEB31111 (14).

#### Pangenome Construction, functional annotation and selection pressure analysis

To quantify the genomic response to fertilization, pangenomes were constructed using Roary (v3.13.0)(15). The distribution of core and accessory genes was calculated to determine the 'Mobile Fortress' capacity of the Bacillus cereus group across different fertilization regimes. Coding sequences (CDS) were predicted from the refined bins and clustered into orthologous groups (gene clusters) using a 95% identity threshold. Functional annotations were assigned to each cluster using the Prokka (v1.14.6) pipeline(16). The resulting gene presence-absence matrix was transformed into a binary format (1 = present, 0 = absent) for downstream statistical enrichment. Next, functional enrichment was determined using two distinct statistical frameworks to distinguish between absolute specialization and relative shifts: absolute enrichment and differential abundance. Absolute enrichment was calculated by hypergeometric distribution test to compare the functional profile of each treatment group against the entire pangenome background and identify statistically over-represented pathways (enrichment score > 1.0; $p < 0.05$) within a specific fertilization regime. To quantify the shift between experimental groups and the control, $\text{Log}_2$ Fold Change (LFC) was calculated for each pathway (differential abundance). The study provided comparative analysis between treatments, identifying fertilization regime-specific gains in metabolic potential. Next, to identify intergenic regions, the reference metagenome previously annotated using Prokka. Genomic features were extracted into GFF3 and BED formats. To determine the spatial context of fixed mutations, we utilized bedtools closest (v2.30.0) to map each intergenic variant to its nearest downstream gene(17). This allowed for the categorization of mutations into three distinct classes: intragenic (within CDS), regulatory (within 0–200 bp of a transcriptional start site), and orphan (deep intergenic, >200 bp from any gene).

To quantify selection pressures within the rhizosphere populations, we calculated ratio of non-synonymous (*dN*) to synonymous (*dS*) substitution rates (ω= *dN/dS*), hereafter referred to as in-situ *dN/dS* to measure in-situ selection coefficient. Because reads were mapped back to de novo assembled consensus MAGs derived from the same sampling event, this metric specifically measures short-term selection intensity and the retention of minority variants within the local population. This approach avoids the consensus bias inherent in metagenomic mapping by focusing on the diversity of the standing genetic variation rather than long-term divergence from an ancestral reference genome. Substitution rates were estimated using the Nei-Gojobori method, which accounts for the proportion of observed substitutions relative to potential synonymous and non-synonymous sites(18). To account for multiple substitutions at a single site (multiple hits), the Jukes-Cantor correction was applied to the raw proportions: $\partial = -\frac{3}{4}\ln\left(1 - \frac{4}{3}p\right)$, where $p$ is uncorrected proportion of *dN* or *dS*. Calculated ω values were used to classify genes into distinct evolutionary regimes —adaptive selection, neutral evolution and purifying selection. The evolutionary hotspot for functional adaption was stated at ω > 1, indicating beneficial amino acid diversification. Neutral evolution (ω ≃ 1) indicates lack of selective constraint; purifying/ conserving selection (the regulatory shield), ω < 1 indicated the removal of deleterious mutations to maintain protein function.

**Community consensus assembly analysis, variant calling and annotation of evolutionary rates**

To understand the evolutionary trajectory of the community, we utilized the pipeline's integrated community consensus assembly (ancestral DNA module) of the nf-core/mag pipeline to reconstruct the ancestral consensus sequences, providing a baseline to differentiate historical lineage traits from recent adaptations. Ancient DNA typically displays characteristic damage, notably cytosine deamination, which appears as C→T transitions. The pipeline uses variant calling to differentiate between true genetic variations and structural damage patterns in the assembled contigs. Phylogenetic placement of metagenome-assembled genomes (MAGs) was performed using GTDB-Tk, followed by the reconstruction of community consensus assembly through the CheckM and BUSCO lineages. This allowed for the identification of core ancestral genes—those conserved across the evolutionary history of the taxa—versus divergent adaptive genes that have emerged or been modified under recent selective pressures. Following assembly, raw reads were re-mapped to the community scaffolds using BWA-MEM aligner(19), FreeBayes (20) in combination with BCFtools (21) was used to call variants from aligned reads. These tools identify positions where the reads differ from the assembled contig, allowing the pipeline to correct errors or re-call the consensus sequence of the contig, effectively mitigating the effects of aDNA damage.

The Single Nucleotide Polymorphisms (SNPs) and small indels were identified using BCFtools (v1.20). Fixed mutation (Allele Frequency (AF) = 1.0) was identified using Bowtie2 in –very-sensitive ensuring that the identified mutations represent traits that have reached 100% saturation within the population, signaling a completed selective sweep(22). To identify potential functional impacts of genomic variations, we integrated variant calls with structural annotations. SNPs and small Indels were stored in VCF format (v4.2), while genome annotations were derived from Prokka (v1.14.6) in GFF3 format. Individual variant call files (VCFs) for each agriculture management regime were indexed and merged using BCFtools. To ensure downstream analytical precision, sample identifiers were mapped to their respective biological treatments (Control,

Organic, and Inorganic) via a custom protocol. This standardized multi-sample VCF served as the primary data source for both the selective sweep detection and the Identity-By-State (IBS) population structure analysis. Global genetic distances were quantified via an Identity-By-State (IBS) framework using 144,465 biallelic SNPs recovered from the metaSPAdes assembly. Due to the high metadata overhead of the metagenomic headers, variant calls were manually extracted from the uncompressed VCF stream and restructured into a diploid-compliant PED format. Missing genotypes were explicitly encoded as null biallelic pairs (0\ 0) to maintain matrix integrity. This high-fidelity approach provided the resolution necessary to distinguish macro-scale genomic drift from localized adaptive selective pressure.

To ensure biological authenticity, raw variants were filtered to remove technical noise common in degraded or low coverage metagenomic data. Specifically, a Subtractive Filtering approach was employed to exclude G ↔ C transversions, which often represent sequencing artifacts rather than true biological variation. The resulting raw VCF files were filtered using BCFtools to retain only high-confidence variants. Filtering thresholds required a minimum read depth (DP) of [10x], a mapping quality (MAPQ) score of [>30], and a minimum Phred base quality (QUAL) of [>30]. Allele Frequencies (AF) were calculated directly from the pooled metagenomic libraries as the ratio of alternate allele observations (AO) to the total mapped read depth at that locus (AF = AO / DP). Fixed mutations were strictly defined as variants reaching an AF of 1.0 (100% alternate allele consensus). Functional annotation of variants was performed by intersecting high-quality SNPs with the Prokka-generated GFF3 files using bedtools v2.30.0. Since the initial variant calls lacked functional metadata, a custom bash pipeline was used to cross-reference SNP coordinates with coding sequences (CDS). Evolutionary rates were estimated by calculating the ratio of non-synonymous to synonymous substitutions ($\omega = dN/dS$). To handle the infinite values typically generated by zero synonymous mutations in small metagenomic windows, a Laplace smoothing factor (pseudocount of 1) was applied to both the numerator and denominator. For all comparative evolutionary metrics, the unit of observation was defined as the individual gene cluster or variant site (n = 5,544 for inorganic; n= 2,038 for organic and n = 2517 for control (unfertilized)). Statistical significance between treatments was determined using the Mann-Whitney U test, which evaluates whether the distribution of evolutionary rates (e.g., $\omega$ or Allele Frequency) differs between populations. This approach treats each gene as an independent outcome of the prevailing selection regime, providing the necessary statistical power to resolve global selection signatures despite the pooled nature of the metagenomic libraries.

*Statistical Analysis and Multivariate Ordination:*

*i) Mutational Signatures and Transition/Transversion (Ti/Tv) Ratios:* Base substitution matrices and global Transition-to-Transversion (Ti/Tv) ratios were extracted directly from the filtered variant files using BCFtools stats. To determine the statistical significance of treatment-induced mutational shifts (e.g., the enrichment of C →T transitions in inorganic soils), expected substitution frequencies were calculated based on the unamended Control baseline. Observed substitution counts were then compared against this expected distribution using a [Binomial Exact Test / Fisher's Exact Test], generating the reported p-values for chemical mutational bias.

*ii) Multivariate Ordination of Selection Pressures:* To classify the adaptive trajectory of the microbiomes, a Principal Component Analysis (PCA) was performed on the *dN/dS* profiles of the annotated genes. Eigenvectors and coordinate matrices were generated via the SPAdes pipeline variants (SPAdes_pca.eigenvec) and plotted using base and multivariate packages in R. To prevent mathematical divide-by-zero errors in genes exhibiting zero synonymous mutations, a standard pseudocount of 1 was applied to both the synonymous and non-synonymous variant counts (pN+1 / pS+1) prior to ordination. The statistical significance of functional gene displacement from the population centroid was validated using Mahalanobis Distance ($D^2$), with genes localizing outside the 95% confidence ellipses ($p < 0.05$) classified as Unique Adaptive Drivers.

**Allele frequency distribution and selection profiling**

To evaluate the intensity of selective sweeps across the three-soil fertilization regime (control, organic, and inorganic), we analyzed the distribution of Allele Frequencies (AF) for all identified variants. To determine if the different fertilization regimes induced distinct selection pressures, we utilized two complementary statistical approaches: (i) *Selection Pressure Distribution:* To test for global shifts in the intensity of selection, we compared the overall AF distributions between the organic and inorganic treatments using a two-sided Mann-Whitney U test. This non-parametric approach was selected due to the characteristic non-normal, skewed distribution of allele frequencies in evolving metagenomes (bimodal peaks identifying selective sweep zone). (ii) *Fixation Proportion Analysis:* To assess whether the mobile genetic elements (MGEs) led to a statistically significant increase in fixed adaptive traits, we performed a Pearson's Chi-squared test on a 2 ×2 contingency table. This table compared the counts of fixed (AF = 1.0) versus non-fixed variants across the contrasting fertilizer treatments.

**Fixation rate and mutation velocity**

We defined the fixation rate as the ratio of fixed mutations to the total number of mutations detected within a sample : $Fixation\ Rate = \frac{Number\ of\ mutation\ with\ AF=1}{Total\ number\ of\ mutations}$. Fixed mutations were identified as those with an allele frequency (AF) of 100%, represented by a genotype of 1/1 (homozygous alternate) in the BCFtools output. The total number of mutations was evaluated using BCFtools, including all variants detected in the samples, comprising both fixed (1/1) and polymorphic/heterozygous (0/1) sites. Mutation counts and fixation rates were integrated into a unified data structure to visualize the relationship between genomic stress (total mutation count) and adaptive success (fixation rate). Statistical significance was defined as $p < 0.05$. All analyses were performed using custom python codes and visualized using Seaborn (v0.12).

**Mutational Velocity and Enrichment Ratios**

Number of fixed mutations per kilobase (Hits/kb) for each contig to allow for comparison across genes of disparate lengths across distinct agriculture regime was measured by quantifying mutational velocity. This metric represents the density of evolutionary hits per 1,000 base pairs of coding sequence. To normalize mutational burden across genes of varying sizes, mutational velocity was calculated as Variants per kilobase (Vpkb) using the explicit formula: $Vpkb = \frac{Total\ variants\ within\ CDS}{Total\ length\ of\ CDS\ in\ base\ pairs} \times 1000$. It defines the fold-change in mutation density of mobile elements relative to this chromosomal baseline. For the non-normal distribution of metagenomic mutation counts, non-parametric statistics were used to test significance. Global variance among treatments was assessed using a Kruskal-Wallis H-test. For pairwise comparisons, we utilized a dual-validation approach to account for unequal sample sizes between the core genome and mobile genetic element (MGE) hotspots. Significance was determined using Mann-Whitney U tests and reinforced with Permutation Tests (10,000 iterations) to provide robust p-values independent of sample size bias. The magnitude of the biological effect (Effect Size) was quantified using Cliff's Delta (d), with thresholds for negligible ($d < 0.147$), small ($d < 0.33$), medium ($d < 0.474$), and large ($d \geq 0.474$) effects. All statistical modeling was performed using custom python codes and statsmodels libraries. Mutation densities were visualized on a $\log_{10}$ scale to accommodate the high-velocity outliers observed in the mobile reservoir. To differentiate targeted regulatory adaptation from stochastic background mutation, an Enrichment Ratio (ER) was calculated for specific functional islands. Background Drift Density was first established by calculating the total number of non-clustered, intergenic variants divided by the total intergenic genomic footprint (in kilobases) across the assembly. The ER was then determined as: $ER = \frac{Local\ density\ \left(\frac{mut}{kb}\right) of\ specific\ region}{Background\ drift\ density\ \left(\frac{mut}{kb}\right)}$.

### Spatial Mapping of Regulatory Mutational Hotspots (spatial genomic shock calculations)

The proximity analysis of intergenic fixed mutations was conducted using bedtools closest with the -d parameter to extract the exact base-pair distance between variants and the nearest annotated coding sequence. A regulatory hit was strictly defined as a variant localizing within the immediate 0–200 bp region upstream of a predicted transcriptional start site. To account for polycistronic operons standard in bacterial genomes, upstream regulatory mutations were assigned to the primary (most proximal) gene in the predicted operon, assuming coordinated downstream transcriptional impact. To establish a statistical baseline for the genomic shock effect, we calculated the mean mutation velocity of the chromosomal compartment (n = X contigs). Fixed variants were intersected with Prokka-annotated features to identify their proximity to coding regions. The promotor region was focused by locating variants positioned within the 0–200 bp upstream of a start codon and were prioritized as potential regulatory adaptations to identify regulatory evolution. For coding-level selection, the non-synonymous to synonymous substitution ratio ($\omega = dN/dS$) for genes were linked to high-density regulatory hotspots. Using a custom python pipeline (incorporating Biopython consensus and translation modules), ($\omega = dN/dS$) values were derived by comparing fixed mutations within the coding sequence (CDS) to the reference sequence. The Enrichment Ratio (ER) was calculated by normalizing the local mutation density of specific functional hubs against the global background of orphan variants (stochastic drift across the intergenic landscape), as given above.

### Mobile Genetic Element (MGE) Identification and genomic partitioning

To partition the metagenome into structural compartments, assembled contigs were processed through geNomad (v1.5.0) in end-to-end mode (23). Contigs were classified into three primary categories based on marker gene content and nucleotide composition: chromosomal, plasmid, and viral. This classification allowed for the stratification of mutational load across mobile and core genomic reservoirs. High-confidence plasmid contigs were isolated based on gene content, amino acid composition, and the presence of plasmid-specific marker genes. To unify the separate annotation and plasmid identification workflows, a custom cross-referencing pipeline was developed. Prokka-generated protein sequences were mapped back to the geNomad plasmid nucleotide databases using blastp and tblastn with a high-stringency threshold (Identity ≥95%, Evalue < 1e-10). Genes successfully mapped to geNomad contigs were partitioned as plasmid-borne, while the remaining genes were categorized as chromosomal.

**Integration of Evolutionary Selection and Mobile Genetic Elements (MGEs) and comparative genomics of highly conserved mobile loci (HCML)**

To determine the genomic partitioning of selective intensity, we developed a multi-omic synthesis pipeline. Gene functional descriptions from Prokka-annotated GFF files were cross-referenced with Metagenome-Assembled Genome (MAG) coordinates to bridge evolutionary selection data ($\omega$= dN/dS) with geNomad-derived mobility scores. High-confidence MGEs were identified using a dual-filter approach: (i) intense purifying selection ($\omega$< 0.1), indicating functional conservation, and (ii) a geNomad mobility score > 0.7, indicating localization to plasmid or viral scaffolds. This integrated workflow allowed for the identification of essential, non-redundant genetic elements within the mobile gene pool of the inorganic and organic rhizospheres. This approach characterized the essential mobile gene pool coined Highly Conserved Mobile Loci (HCML) (coding sequences (CDS) that are localized on mobile genetic elements (geNomad score > 0.99 and/or recovery in the metaSPAdes --metaplasmid track) and exhibit evidence of intense purifying selection ($\omega$= dN/dS < 0.1).

To refine the candidate list of HCMLs, a post-processing script was implemented to resolve data redundancies and prioritize biologically relevant features. The initial multi-omic merge often generated inflated record counts (e.g., >400,000 rows) due to the combinatorial matching of non-unique functional descriptions, such as hypothetical protein. To correct this, a spatial deduplication protocol was applied. The dataframes were combined based on unique pairs of scaffold ID and functional annotation, ensuring that each discrete gene locus on a predicted mobile element was represented as a single entry, regardless of how many times that specific annotation appeared across the global dataset. A selective column filter was applied to retain only the most critical parameters for comparative genomics which include selective pressure, geNomad-derived plasmid and virus confidence scores and standardized gene product description. Final HCML tables were sorted in ascending order of *dN/dS* ratio. This prioritization highlights genes under the most intense purifying selection (*dN/dS* → 0), identifying the evolutionary foundation of the soil mobilome—genes that are essential for survival under the selective pressures of the respective nitrogen treatments. Protein sequences for all identified HCMLs were extracted from Prokka-annotated amino acid files (.faa) using custom python scripts. To assess the divergence of the conserved mobilome between fertilization regimes, we performed global sequence clustering using CD-HIT (v4.8.1) with a 95% identity threshold and a word size of 5 (c=0.95, n=5). This allowed for the categorization of HCMLs into treatment-unique clusters and a shared core soil mobilome.

**Physical Characterization of the Cryptic Mobilome**

Following the identification and strict curation of Highly Conserved Mobilome Lineages (HCMLs) unique to each nitrogen regime, the full-length genomic contigs harboring these loci were computationally isolated from the bulk metagenomic assemblies. A custom Python processing pipeline utilizing the Biopython library (v1.81) was deployed to parse the original SPAdes-generated assembly files (.fasta.gz). Contigs mapping exactly to the filtered HCML NODE_ID identifiers were extracted into treatment-specific FASTA files, yielding 122 genomic contigs for the inorganic treatment and 75 contigs for the organic treatment for downstream architectural profiling.

To elucidate the physical properties and horizontal transfer potential of these genetic vehicles, the extracted sequences were analyzed using mob_typer from the MOB-suite software package(24). The tool was executed in multi-sequence mode (--multi) to generate detailed, per-contig architectural profiles. Core physical metrics, including total sequence length, GC content (%), and topological circularity, were recorded. The topological circularity of the contigs was utilized as a primary proxy for identifying intact, circular episomal elements or cryptic plasmids. Furthermore, mob_typer was used to annotate mobility-associated genetic features, specifically screening for replicon types, mate-pair formation (MPF) complexes (e.g., Type IV secretion systems), origins of transfer (oriT), and relaxase protein families (e.g., MOBP, MOBV). Based on the presence or absence of these mobility signatures, each contig was functionally categorized regarding its horizontal transfer potential (conjugative, mobilizable, or non-mobilizable). The putative biological sources and host ranges of the sequestered HCML-carrying contigs were determined using the MASH-based nearest-neighbor identification algorithm integrated within mob_typer (25). Each sequence was queried against the comprehensive MOB-suite reference database to assign the closest taxonomic neighbor. This enabled the comparative assessment of host-range dynamics and taxonomic shifts within the mobilome driven by the distinct fertilization strategies.

**RNA-Seq Data Analysis:** Secondary analysis of bulk RNA-seq (26) FASTQ files was performed with the nf-core/rnaseq pipeline. The analysis includes initial QC, trimming, alignment with reference genome “GCF_902167145.1”, gene quantification, and post-alignment QC. The tertiary data analysis with additional QC, normalization, and differential gene expression analysis between pairwise comparison was performed with the DESeq2 package(27). Significantly expressed DEGs were defined by ($\log2FC > |2|$, $P_{adj} < 0.05$). Further, gene enrichment analysis was performed with FGSEA(28).

**Network Construction:** The regulator-pathway and shared host-microbe interaction networks were generated in Python from integrated host transcriptomic, GO enrichment, and microbial mutation-annotation datasets for the inorganic treatment comparison. For the regulator-pathway network, differentially expressed maize genes annotated as regulators were extracted from the inorganic-versus-control DESeq2 results and linked to significantly enriched GO terms when the regulator gene appeared in the term-specific gene membership list. The regulator nodes were labeled using host annotation metadata, colored by direction of differential expression, and connected to GO nodes scaled by term gene count and colored by significantly up or down regulation. For the shared host-microbe network, overlapping GO terms between host and microbial datasets were used to connect differentially expressed host genes to microbial proteins associated with mutation-supported enriched pathways. Host genes were required to occur in at least two shared GO terms, while microbial seed nodes were defined by mutated proteins and expanded to same-operon neighboring coding sequences identified from the metagenomic GFF annotation on the same contig and strand with intergenic gaps of at most 100 bp. The resulting network displayed shared GO pathways as central nodes, with host nodes colored by transcriptional response and microbial nodes distinguished as directly mutated or downstream operon-associated proteins.

**Statistical Analysis:** All statistical integrations and visual representations of the mobilome architecture were performed using Python (v3.x) with the pandas, seaborn, and matplotlib libraries (29, 30). To determine whether the contrasting nitrogen regimes induced significant structural shifts in the mobilome, quantitative comparisons of continuous genomic variables—specifically contig size (bp) and GC content (%)—were conducted. Because these genomic features often exhibit non-normal distributions, the non-parametric Mann-Whitney U test was employed via the scipy.stats module. Differences in the categorical distribution of mobility potential (mobilizable vs. non-mobilizable) between the two treatments were evaluated using a Chi-squared test of independence. For all analyses, a p-value threshold of $< 0.05$ was established to indicate statistical significance. All statistical analysis and visualization for RNA-seq was performed in the R programming language.


1. O. O. Babalola, R. R. Molefe, A. E. Amoo, Metagenome Assembly and Metagenome-Assembled Genome Sequences from the Rhizosphere of Maize Plants in Mafikeng, South Africa. *Microbiology Resource Announcements* **10**, e00954-00920 (2021).
2. S. Andrews (2010) FastQC: A Quality Control Tool for High Throughput Sequence Data.
3. P. Ewels, M. Magnusson, S. Lundin, M. Käller, MultiQC: summarize analysis results for multiple tools and samples in a single report. *Bioinformatics* **32**, 3047-3048 (2016).
4. P. A. Ewels *et al.*, The nf-core framework for community-curated bioinformatics pipelines. *Nature Biotechnology* **38**, 276-278 (2020).
5. S. Krakau, D. Straub, H. Gourlé, G. Gabernet, S. Nahnsen, nf-core/mag: a best-practice pipeline for metagenome hybrid assembly and binning. *Nar Genom Bioinform* **4** (2022).
6. B. Bushnell (2014) BBMap: A Fast, Accurate, Splice-Aware Aligner. (Lawrence Berkeley National Laboratory (LBNL), Berkeley, CA (United States)).
7. D. Kim, L. Song, F. P. Breitwieser, S. L. Salzberg, Centrifuge: rapid and sensitive classification of metagenomic sequences. *Genome Research* **26**, 1721-1729 (2016).
8. S. Nurk, D. Meleshko, A. Korobeynikov, P. A. Pevzner, metaSPAdes: a new versatile metagenomic assembler. *Genome Research* **27**, 824-834 (2017).
9. J. Alneberg *et al.*, Binning metagenomic contigs by coverage and composition. *Nature Methods* **11**, 1144-1146 (2014).
10. D. D. Kang *et al.*, MetaBAT 2: an adaptive binning algorithm for robust and efficient genome reconstruction from metagenome assemblies. *PeerJ* **7**, e7359 (2019).
11. C. M. Sieber *et al.*, Recovery of genomes from metagenomes via a DAS Tool-driven consensus approach. *Nature Microbiology* **3**, 836-843 (2018).
12. M. Manni, M. R. Berkeley, M. Seppey, F. A. Simao, E. M. Zdobnov, BUSCO Update: Novel and Streamlined Workflows along with Broader and Deeper Phylogenetic Coverage for Scoring of Eukaryotic, Prokaryotic, and Viral Genomes. *Mol Biol Evol* **38**, 4647-4654 (2021).
13. D. H. Parks, M. Imelfort, C. T. Skennerton, P. Hugenholtz, G. W. Tyson, CheckM: assessing the quality of microbial genomes recovered from isolates, single cells, and metagenomes. *Genome Res* **25**, 1043-1055 (2015).
14. J. Nelkner *et al.*, Effect of Long-Term Farming Practices on Agricultural Soil Microbiome Members Represented by Metagenomically Assembled Genomes (MAGs) and Their Predicted Plant-Beneficial Genes. *Genes (Basel)* **10** (2019).

15. F. Sitto, F. U. Battistuzzi, Estimating Pangenomes with Roary. *Mol Biol Evol* **37**, 933-939 (2020).
16. T. Seemann, Prokka: rapid prokaryotic genome annotation. *Bioinformatics* **30**, 2068-2069 (2014).
17. A. R. Quinlan, BEDTools: The Swiss-Army Tool for Genome Feature Analysis. *Curr Protoc Bioinformatics* **47**, 11 12 11-34 (2014).
18. M. Nei, T. Gojobori, Simple methods for estimating the numbers of synonymous and nonsynonymous nucleotide substitutions. *Mol Biol Evol* **3**, 418-426 (1986).
19. H. Li, Aligning sequence reads, clone sequences and assembly contigs with BWA-MEM. *arXiv preprint arXiv:1303.3997* (2013).
20. E. Garrison, G. Marth, Haplotype-based variant detection from short-read sequencing. *arXiv preprint arXiv:1207.3907* (2012).
21. P. Danecek *et al.*, Twelve years of SAMtools and BCFtools. *Gigascience* **10** (2021).
22. B. Langmead, S. L. Salzberg, Fast gapped-read alignment with Bowtie 2. *Nat Methods* **9**, 357-359 (2012).
23. L. Bar, R. J. Darrah, S. Vaidyanathan, Analysis of the Genome Aggregation Database (gnomAD) reveals a global burden of cystic fibrosis and the need for improved diagnosis and care. *EBioMedicine* **124**, 106145 (2026).
24. J. Robertson, J. H. Nash, MOB-suite: software tools for clustering, reconstruction and typing of plasmids from draft assemblies. *Microbial Genomics* **4**, e000206 (2018).
25. K. Abram *et al.*, Mash-based analyses of Escherichia coli genomes reveal 14 distinct phylogroups. *Commun Biol* **4**, 117 (2021).
26. B. W. Luo *et al.*, Fertilization regulates maize nutrient use efficiency through soil rhizosphere biological network and root transcriptome. *Appl Soil Ecol* **207** (2025).
27. M. I. Love, W. Huber, S. Anders, Moderated estimation of fold change and dispersion for RNA-seq data with DESeq2. *Genome Biol* **15**, 550 (2014).
28. G. Korotkevich *et al.*, Fast gene set enrichment analysis. *bioRxiv* (2021).
29. P. Virtanen *et al.*, SciPy 1.0: fundamental algorithms for scientific computing in Python. *Nature Methods* **17**, 261-272 (2020).
30. M. L. Waskom, Seaborn: statistical data visualization. *Journal of Open Source Software* **6**, 3021 (2021).
